# The mitochondrial-targeted peptide therapeutic elamipretide improves cardiac and skeletal muscle function during aging without detectable changes in tissue epigenetic or transcriptomic age

**DOI:** 10.1101/2024.10.30.620676

**Authors:** Wayne Mitchell, Gavin Pharaoh, Alexander Tyshkovskiy, Matthew Campbell, David J. Marcinek, Vadim N. Gladyshev

## Abstract

Aging-related decreases in cardiac and skeletal muscle function are strongly associated with various comorbidities. Elamipretide (ELAM), a novel mitochondrial-targeted peptide, has demonstrated broad therapeutic efficacy in ameliorating disease conditions associated with mitochondrial dysfunction across both clinical and pre-clinical models. ELAM is proposed to restore mitochondrial bioenergetic function by stabilizing inner membrane structure and increasing oxidative phosphorylation coupling and efficiency. Although ELAM treatment effectively attenuates physiological declines in multiple tissues in rodent aging models, it remains unclear whether these functional improvements correlate with favorable changes in molecular biomarkers of aging. Herein, we investigated the impact of 8-week ELAM treatment on pre- and post-measures of C57BL/6J mice frailty, skeletal muscle, and cardiac muscle function, coupled with post-treatment assessments of biological age and affected molecular pathways. We found that health status, as measured by frailty index, cardiac strain, diastolic function, and skeletal muscle force are significantly diminished with age, with skeletal muscle force changing in a sex-dependent manner. Conversely, ELAM mitigated frailty accumulation and was able to partially reverse these declines, as evidenced by treatment-induced increases in cardiac strain and muscle fatigue resistance. Despite these improvements, we did not detect statistically significant changes in gene expression or DNA methylation profiles indicative of molecular reorganization or reduced biological age in most ELAM-treated groups. However, pathway analyses revealed that ELAM treatment showed pro-longevity shifts in gene expression such as upregulation of genes involved in fatty acid metabolism, mitochondrial translation and oxidative phosphorylation, and downregulation of inflammation. Together, these results indicate that ELAM treatment is effective at mitigating signs of sarcopenia and heart failure in an aging mouse model, but that these functional improvements occur independently of detectable changes in epigenetic and transcriptomic age. Thus, some age-related changes in function may be uncoupled from changes in molecular biological age.

## INTRODUCTION

Given the rapidly aging world population, it is expected that healthcare costs associated with treating cancer, cardiovascular disease, and certain neurological conditions will surge in the coming decades^1^. According to the geroscience hypothesis, interventions that target the underlying physiology of the aging process would represent the most utilitarian modality to mitigate aging-related declines in physical/functional health across multiple systems. Thus, there is an unmet need for safe and well-tolerated geroprotectors that can lead to significant improvements in overall health during aging. In particular, sarcopenia affects up to 45% of the senior population^2^, and aging-related heart failure is the major cause of mortality in older adults^3,4^. The Interventions Testing Program (ITP) has successfully identified a number of geroprotectors that are able to extend healthspan and lifespan in genetically diverse mice^5–7^; in turn, some of these drugs may also have beneficial effects on muscle performance^8,9^. However, a consistent trend with these studies has been that the studied compounds are much more likely to improve healthspan and lifespan in males than in females^6,7,10^. Moreover, many interventions that extend lifespan do not strictly act on aging-specific features but rather alter the signaling of developmental and/or growth-related pathways^11,12^. Therefore, it is imperative to develop next-generation drug candidates that directly target aging-related molecular features and translate equally as well across the sexual dimorphism of human aging^13^.

Elamipretide (ELAM) is a novel mitochondrial-targeted therapeutic peptide, currently under clinical investigation, that has demonstrated broad scope efficacy in treating a wide array of pathologies associated with mitochondrial dysfunction. This includes rare genetic diseases such as Friedreich’s ataxia^14–16^ and Barth syndrome^17–20^, as well as conditions normally associated with aging such as ischemia-reperfusion injury^21–24^, neurological disorders^25–27^, dry age-related macular degeneration^28–30^, and heart failure^31,32^. ELAM is known to concentrate up to 1,000-fold in mitochondria and associate with the cardiolipin-rich inner mitochondrial membrane^23,24^, while simultaneously demonstrating minimal toxicity and favorable pharmacokinetics^17,28,33–35^. Several non-mutually exclusive mechanisms have been proposed to explain their ability to improve mitochondrial bioenergetic function in various conditions of cellular stress and disease. These include interactions with mitochondrial proteins involved in energy generation^36^, reducing proton leak through the adenine nucleotide translocase (ANT) protein^31,37,38^, down-tuning inner membrane surface electrostatics to reduce/alter mitochondrial Ca^2+^ influx^39,40^, and modulating inner membrane biophysical properties^39^. In support of the latter two mechanisms, a recent study has shown that the ability of this class of peptides to attenuate inner membrane surface electrostatics is well-correlated with their ability to improve ATP production and cell viability^40^.

While the functional improvements in aging models following ELAM treatment are well-documented^38,41–43^, it is unknown whether they strongly correlate with favorable changes in molecular biomarkers of aging. Biomarkers of aging are increasingly being validated for their ability to predict disease and aging-related health outcomes^44,45^. The latest epigenetic clocks can predict mortality^46,47^, distinguish adaptive and causal changes in DNA methylation (DNAm)^48^, and utilize universal gains in methylation at Polycomb Repressive Complex 2 target regions to predict cellular age^49^. Additionally, plasma-based proteomics clocks are now able to identify age acceleration in specific organs^50,51^, and recently developed multi-tissue transcriptomic clocks can predict the effects of lifespan-shortening and lifespan-extending interventions and provide insights into the specific molecular processes involved in organismal aging and mortality^52^. However, concerns have been raised about the potential clinical use of some of these biomarkers, particularly in regards to their interpretability^53,54^, their ability to distinguish inflammation from aging^55,56^, and whether or not they successfully report on improvements in tissue and organ function^57,58^.

In this work, to evaluate the correlation between functional changes in muscle performance with biomolecular features of aging, we administered ELAM for 8 weeks to young (5-month-old) and old (24-month-old), male and female C57BL/6J mice. We performed both pre- and post-treatment measures of frailty index (FI), *in vivo* muscle force, and cardiac function using echocardiography, which allowed for longitudinal assessment of the effects of ELAM treatment. At the end of the study, RNA isolated from cardiac and skeletal muscle tissues were submitted for bulk mRNA-seq; in addition, the effect of ELAM on DNAm in cardiac tissue was determined by DNAm microarray. Using the obtained mRNA-seq and DNAm data, we examined cross-sectional changes in molecular organization induced by ELAM treatment along with its effect on biological age estimated with transcriptomic and epigenetic clocks, and on gene expression signatures of aging, mortality, and longevity^52,59,60^. In agreement with previous studies^38,41–43^, we found that cardiac systolic and diastolic function deteriorates during aging and that treatment with ELAM can mitigate these effects, as made evident by significant improvements in global longitudinal strain and ejection fraction, as well as reduced frailty accumulation in aged mice. Moreover, we discovered that muscle force across a range of frequencies and fatigue stimulations decreases with aging, particularly in female mice. In turn, chronic ELAM treatment was able to attenuate this decline in female mice. While transcriptomic changes caused by ELAM treatment showed significant positive correlation with signatures of mammalian longevity, no significant effect on the expression of individual genes, methylation status of CpG sites, and overall epigenetic or transcriptomic age was detected for most treatment groups. Taken together, the results of our study highlight that improvements in tissue function during aging can occur without substantial changes in molecular profiles and associated biological age. In addition, our study further identifies ELAM as a novel geroprotector that demonstrates both age-specific effects and an equivalent ability to improve healthspan in males and females.

## MATERIALS AND METHODS

### Animals

This study was reviewed and approved by the University of Washington Institutional Animal Care and Use Committee (IACUC). Male and female C57BL/6J mice were received from the National Institute on Aging mouse colony and housed at the University of Washington in a specific-pathogen free facility. All mice were maintained at 21°C on a 14/10 light/dark cycle at 30-70% humidity and given standard mouse chow (LabDiet PicoLab® Rodent Diet 20) and water *ad libitum*.

### Experimental design and ELAM treatment

Male and female mice underwent baseline frailty index, *in vivo* muscle force, and echocardiography at 4-5 months in young mice and 23-24 months in old mice. ELAM was delivered to mice for 8 weeks as previously described using subcutaneously (SQ) implanted osmotic minipumps^61^. ELAM treatment was initiated at 5 months (young) and 24 months (old). Midpoint frailty index was measured after 4 weeks, and ELAM pumps were surgically replaced. Post-treatment frailty index, *in vivo* muscle force, and echocardiography were performed between 6-8 weeks of ELAM treatment. Animals were euthanized and tissues (gastrocnemius muscle, heart) were collected at 7 months (young) and at 26 months (old). Tissues of four to five animals per group were used for mRNA-seq and DNAm processing and analysis.

### Frailty index

A 31-item frailty index was assessed longitudinally by the same researcher (G.P.) as previously described^62^. Body mass and temperature scoring were determined using standard deviations from the mean of young or old control mice at baseline.

### In vivo assessment of cardiac function and skeletal muscle force

Echocardiography was performed as previously described with some modifications^38^. In addition to the previous method quantifying parasternal long axis view (PLAX) B-mode and parasternal short-axis view (PSAX) B-mode and M-mode images, pulse wave doppler and tissue doppler 4 chamber view images were quantified to measure diastolic function. *In vivo* measurement of muscle force was performed as previously described in the hindlimb plantar flexor muscles^63^.

### mRNA-seq

RNA, DNA, and protein were isolated from ∼30 mg of flash-frozen cardiac and gastrocnemius tissue. The tissues were homogenized in 2 ml tubes pre-filled with 2.8 mm ceramic beads on a Bead Ruptor Elite tissue homogenizer (OMNI International, Kennesaw, GA). Total RNA, DNA, and protein were purified from the homogenized tissues using the Allprep Mini kit according to the manufacturer’s instructions (Qiagen, Hilden, Germany). RNA concentration was measured by Qubit using the RNA HS assay (ThermoFisher, Waltham, MA). Libraries were prepared using poly A enrichment according to procedures described previously^64^ and sequenced with Illumina NovaSeq X Plus (PE150). Fastq files were mapped to the mm10 (GRCm38.p6) mouse genome using STAR v2.7.2b^65^.

Raw expression data was subjected to filtering, and genes with at least 10 reads in 12.5% of samples were kept for subsequent analysis. Gene expression profiles were subjected to RLE normalization, log transformation, and YuGene normalization, and centered around median expression profiles of young male controls. Differentially expressed genes were identified using custom models in R with DESeq2 3.13 and edgeR 3.34.1^66^. The missing values corresponding to clock genes not detected in the data were imputed with the precalculated average values. Relative transcriptomic age (tAge) of mouse tissues were predicted using the mouse multi-tissue transcriptomic clocks of chronological age and expected mortality based on Bayesian Ridge models^52^. Pairwise differences between average tAges were assessed with mixed-effect models. Resulting p-values were adjusted with the Benjamini-Hochberg (BH) method. Module-specific transcriptomic clocks were applied to scaled relative gene expression profiles in a similar manner. Differences in average transcriptomic ages between age- and sex-matched control and ELAM-treated samples were assessed using a t-test and normalized by dividing the standard deviation of tAge differences across clocks. BH method was used to adjust p-values from modular clock analysis for multiple comparisons.

Gene set enrichment and Spearman correlation analyses against previously established gene expression signatures of mammalian aging, mortality, maximum lifespan, and specific lifespan-extending interventions (rapamycin, caloric restriction, growth hormone deficiency) were performed as described^52,59,64^. GSEA was conducted on pre-ranked lists of genes based on log_10_ (p-value) corrected by the sign of regulation, calculated as: log (pv) x sgn (lfc), where pv and lfc are the p-value and logFC of a certain gene, respectively, obtained from edgeR output, and sgn is the signum function (equal to 1, −1, and 0 if value is positive, negative, or equal to 0, respectively). HALLMARK, KEGG, and REACTOME ontologies from the Molecular Signature Database (MSigDB) were used as gene sets for GSEA. The GSEA algorithm was performed separately for each signature via the fgsea package in R with multilevel splitting Monte Carlo approach and 5000 permutations. P-values were adjusted with BH method. Gene Ontology (GO) terms significantly affected by ELAM treatment (BH adjusted p-values < 0.05) were reduced using REVIGO^67^ and visualized using Cytoscape^68^.

### DNAm

Genomic DNA concentration was measured using the Qubit dsDNA HS assay kit (ThermoFisher, Waltham, MA). DNAm microarray data was obtained on the Horvath mammal 320k array^69^ through the Epigenetic Clock Development Foundation (Torrance, CA). The raw methylation array data was normalized using SeSAMe^70^ and analyzed in the R/Bioconductor environment using methods previously described^64^.

### Western blot

The dried protein pellets obtained from the Allprep purifications were solubilized in 5% SDS for ∼2 hours on a rotisserie at room temperature. Protein concentration was measured using the Pierce BCA kit (ThermoFisher, Waltham, MA) in clear 96-well plates and the absorbance of each sample at 532 nm was measured on a BioTek Synergy H1 microplate reader (Winooski, VT). Samples were diluted to 2 mg/ml in 5% SDS and 1X SDS-PAGE sample buffer (+ 50 mM TCEP), and 30 µg of each sample were loaded and run on Mini-PROTEAN TGX 4-15% acrylamide gels (Bio-Rad, Hercules, CA). Proteins were transferred to PVDF membranes, blocked with 10% dry milk in TBS-T, and incubated at 4°C overnight with the following antibodies (all at a dilution factor of 1:1000): mouse anti-β-actin (Santa-Cruz Biotechnology, Dallas, TX) and rabbit anti-TFAM, anti-MGMT, and anti-Hsp70 (Cell Signaling Technology, Danvers, MA). After washing twice with TBS-T, membranes were incubated with either anti-mouse or anti-rabbit IRDye 680RD (LI-COR Biosciences, Lincoln, NE) for 2 hours at room temperature before washing, drying, and imaging on an Odyssey Fc Imager (LI-COR Biosciences, Lincoln, NE). Band intensities were quantified in FIJI^71^ and TFAM, Hsp70, and MGMT abundances were normalized to the values obtained from the respective β-actin bands.

### Statistical analysis

Graphing and statistical analysis for frailty index, muscle force, and echocardiography data were performed using Microsoft Office Excel and GraphPad Prism for OS X (GraphPad Software, San Diego, California USA). For all statistical tests, alpha levels were set to p < 0.05. *p < 0.05 for post hoc tests or direct comparisons. All statistical tests performed are indicated in the respective figure legends, and exact p-values are depicted in the figures themselves. For all omics experiments, a minimum of n=4 independent biological replicates per treatment condition were utilized. For Western blots, n=3 independent biological replicates per treatment condition were used.

## RESULTS

### Study design

To randomize animals for subsequent analyses, young (4-5-month-old) and old (23-24-month-old) male and female C57BL/6J mice underwent baseline frailty index, *in vivo* muscle force, and echocardiography measurements (Figure 1A). Then, ELAM (3 mg/kg/day) or vehicle (isotonic saline) were delivered to the mice for 8 weeks using two consecutive SQ implanted micro-osmotic pumps, and post-treatment frailty index, *in vivo* muscle force, and echocardiography measurements were performed after the second pump was explanted. Following completion of post-treatment testing, gastrocnemius muscle and heart samples were collected for the cross-sectional molecular analyses.

**Figure 1:**
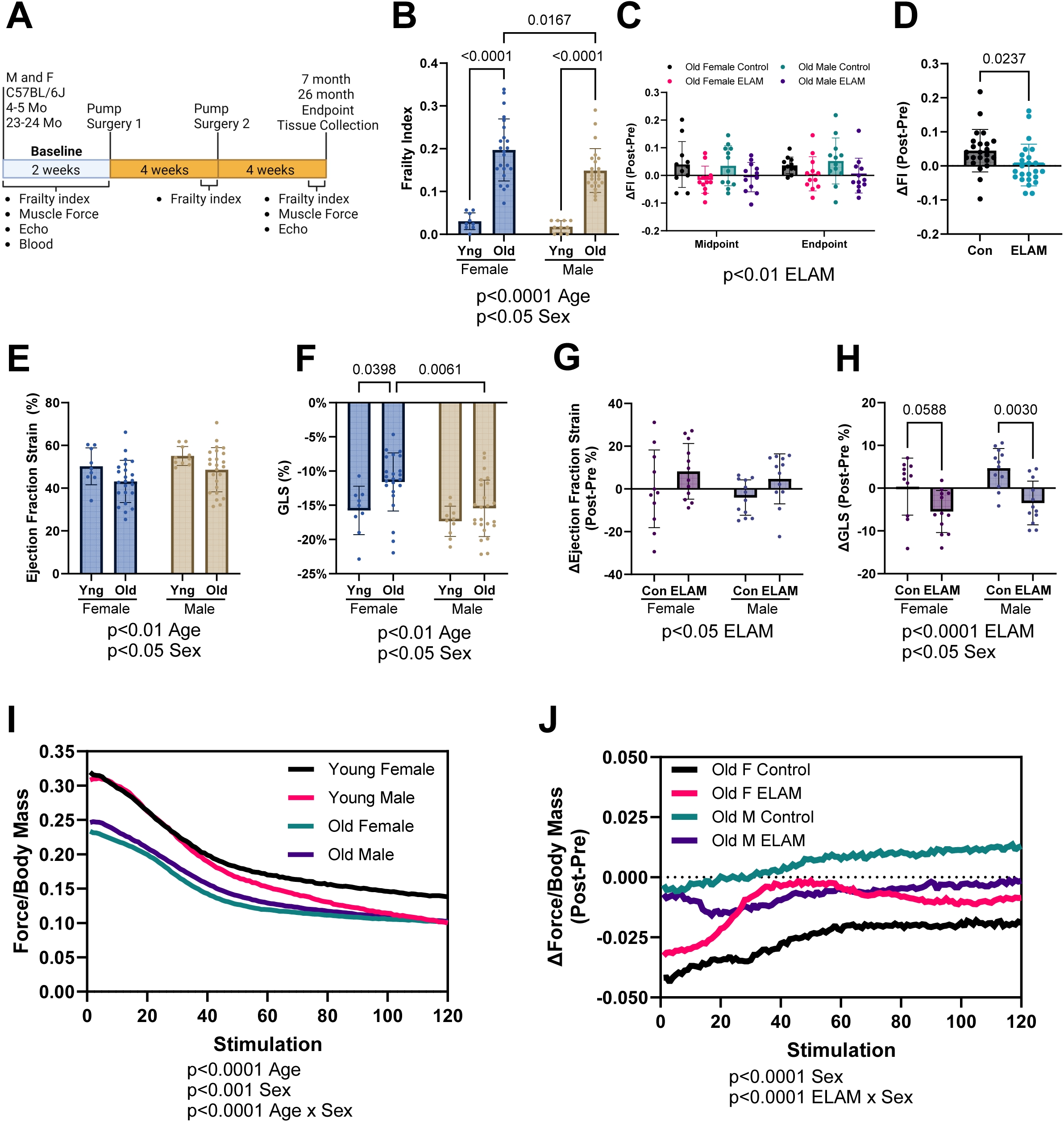
Effect of 2-months ELAM treatment on frailty, cardiac function, and muscle force. A. Experimental design. B. Baseline frailty index (FI) of young (Yng) and old mice. C. Delta (Δ, Post-pre) FI at midpoint (4 weeks) and endpoint (8 weeks) for old mice. D. Combined Δ frailty index at endpoint for control (Con) and elamipretide (ELAM) treated male and female old mice. E. Baseline ejection fraction determined and F. Global longitudinal strain (GLS) from strain analysis of parasternal long axis images of the left ventricle (LV) of young and old mice. G. Δ ejection fraction and H. GLS determined using strain analysis at endpoint for old mice. I. Baseline in vivo muscle force normalized to body mass during repeating fatigue stimulations in young and old mice. J. Δ in vivo muscle force normalized to body mass during repeating fatigue stimulations at endpoint for old mice. 4-5-month-old (young) female and male (n=9-10) and 23-24-month-old (old) female (n=23-24) and male were compared for each measurement at baseline. ELAM treatment effects were compared in control and ELAM-treated old mice (n=11-12) using Δ measurements. Only mice that survived to study endpoint were included in the analysis. Statistical significance was determined by two-way ANOVA with Tukey’s post-hoc test, except for *in vivo* muscle fatigue and ΔFI which were determined by three-way ANOVA. Combined ΔFI significance determined by student’s t-test. Significant ANOVA factors written in text with selected Tukey’s post hoc test comparisons on graphs. Error bars represent sample means ± standard deviations; error bars omitted from I., J. for clarity.

### ELAM treatment mitigates frailty accumulation in aged mice

We used a 31-point frailty score to assess the aging trajectories of the mice, as increased frailty is predictive of adverse health outcomes^62^. Frailty index was significantly increased in aged male and female mice at baseline, with aged female mice exhibiting slightly higher frailty than aged males (Figure 1B). Increased age-related frailty was caused by the accumulation of similar index deficits in old male and female mice (Extended Data Figure 1). Old control mice continued to accumulate frailty scores during the 8-week treatment period, whereas ELAM treatment significantly blunted this increase in frailty in both sexes (Figure 1C-D). Thus, we concluded that ELAM administration can mitigate frailty progression in both old male and female mice.

### ELAM treatment improves cardiac and skeletal muscle function in old mice

Next, we assessed the longitudinal impact of age and ELAM treatment on several parameters related to cardiac and skeletal muscle function. From our echocardiography experiments, we observed that cardiac function was significantly impaired with age in both sexes. Using strain analysis, we identified age-related reduced ejection fraction, fractional shortening, impaired global longitudinal strain (GLS), and cardiac hypertrophy (Figure 1E-F, Extended Data Figure 2A-H). Moreover, reduced GLS and ejection fraction are known predictors of heart failure and mortality in humans and mice^72–76^. Importantly, treatment with ELAM improved these measures of cardiac function in aged male and female mice, although there was no effect on fractional shortening or cardiac hypertrophy (Figure 1G-H, Extended Data Figure 2I-P). Conventional echocardiography identified larger diameter and volumes of the left ventricle (LV) with age (Extended Data Figure 3A-H). Diastolic function was mildly impaired in aged mice, as measured by decreased E’/A’ (Extended Data Figure 4A-H). Aged mice demonstrated cardiac hypertrophy including increased LV wall thickness, LV mass, and heart mass (Extended Data Figure 5A-H). However, these measures were not significantly impacted by ELAM treatment in this study (Extended Data Figure 3I-P, 4I-P, 5I-P). Therefore, we concluded that chronic ELAM administration can partially reverse age-related signs of heart failure in mice mainly by improving systolic function.

In tandem, we measured *in vivo* muscle force production in the hindlimb muscles using force-frequency and fatiguing stimulation protocols. Gastrocnemius muscle mass decreased with age and was not affected by ELAM treatment (Extended Data Figure 6A-B). Across the range of increasing stimulation frequencies used, muscle force was significantly reduced in both aged male and female mice (Extended Data Figure 6C), with the age-related decline in skeletal muscle performance being more pronounced in female mice. Additionally, maximum force production was strongly reduced with age (Extended Data Figure 6D). ELAM treatment did not improve force production in the force frequency protocol or maximum force production (Extended Data Figure 6E-F). Muscle contraction and relaxation speeds were also decreased by age; in contrast, ELAM improved muscle relaxation speeds in male mice (Extended Data Figure 6G-J). In the fatiguing protocol, the hindlimb muscles of aged mice suffered from greater fatigue (Figure 1I), with aged female mice showing more pronounced declines than aged males. While ELAM treatment did not improve muscle force in the aged animals using force-frequency and maximum force production protocols, it did mitigate losses in force production in the fatiguing protocol for aged female mice (Figure 1J). In the fatigue protocol, contraction and relaxation speeds also decreased with age, and ELAM treatment resulted in improved muscle relaxation speeds in aged male mice (Extended Data Figure 7A-D). Taken together, we determined that ELAM affected skeletal muscle function in aged mice, including improved muscle relaxation speeds and maintaining muscle force during fatiguing stimulation.

### ELAM treatment upregulates pathways related to mitochondrial function and energy metabolism

To gain insights into the molecular pathways affected by ELAM treatment, we performed mRNA-seq on transcripts expressed in cardiac and skeletal muscle tissues following completion of post-treatment functional tests. By principal components analysis (PCA) (Figure 2A), we observed clustering of samples primarily by age in both skeletal muscle (left) and heart (right) tissues. When we performed differential expression analysis, most treatment conditions resulted in zero differentially expressed genes (adjusted p-value < 0.05) with the exception of young male and female hearts (Figure 2B). However, at the level of enriched pathways, many aging-associated biological processes were affected by ELAM treatment (Figure 2C). Across both sexes and age groups, we observed significant enrichment of upregulated genes in gene sets associated with oxidative phosphorylation (OXPHOS), mitochondrial translation, and fatty acid metabolism, which resembled the effect of rapamycin treatment and was positively correlated with the functional signature of rodent maximum lifespan. Importantly, expression of these pathways is strongly suppressed during aging, and upregulated mitochondrial translation is a shared mechanism of longevity both within and across mammalian species^60^. In contrast, most pathways downregulated by ELAM treatment such as TNF-ɑ signaling and interferon gamma response are associated with immune-related processes that undergo increased expression during aging. Interestingly, these effects were only not observed in young male and female skeletal muscle, which clustered separately away from the other experimental samples. Consistently, gene expression changes induced by ELAM in all cases except for skeletal muscle in young females were positively correlated with signatures of lifespan and rapamycin, and negatively correlated with most signatures of aging and mortality, both at the level of individual genes (Figure 2D) and enriched pathways (Extended Data Figure 8A). Thus, we concluded that although ELAM treatment did not induce significant changes in gene expression profiles of murine heart and skeletal muscle, the direction of its effect resembled that of rapamycin, one of the most robust and validated interventions^77^. Moreover, ELAM treatment positively affected molecular pathways both strongly associated with aging and longevity, consistent with ELAM’s proposed mechanisms^36,39^.

**Figure 2:**
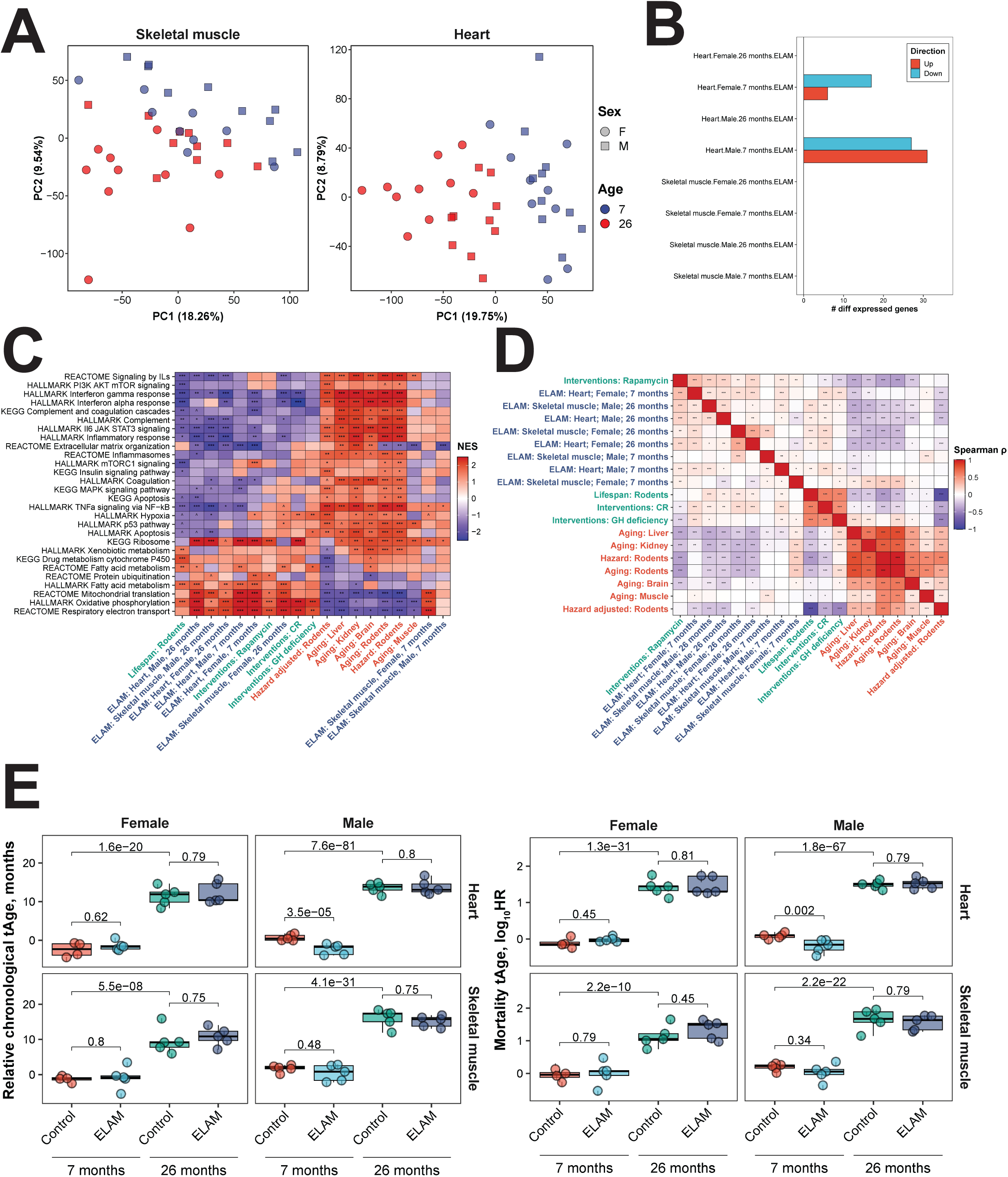
Effect of 2-months ELAM treatment on the transcriptome of young and old mouse heart and skeletal muscle tissues. *A. Principal components analysis (PCA) of mRNA-seq samples.* PCA was performed following filtering of genes with low numbers of reads and RLE normalization^64^. *B. Number of differentially expressed genes following ELAM treatment.* Differentially expressed genes were determined using edgeR^66^ separately for males and females. P-values were adjusted for multiple comparisons using the Benjamini-Hochberg method, and the False Discovery Rate (FDR) was set at 5%. *C. Gene set enrichment analysis (GSEA) of pathways affected by signatures of aging, mortality, lifespan-extending interventions, and ELAM treatment.* Normalized enrichment scores (NES) of aging-related pathways affected by ELAM treatment (blue), lifespan-increasing interventions (green), and signatures of aging and mortality (red). GH: grown hormone deficiency, CR: caloric restriction. ‘adjusted p-value < 0.1, *adjusted p-value < 0.05, **adjusted p-value < 0.01, ***adjusted p-value < 0.001. *D. Correlation analysis of gene expression changes induced by ELAM treatment at the level of enriched pathways.* Spearman correlation between normalized enrichment scores (NES) from gene set enrichment analysis (GSEA) performed for signatures of ELAM treatment (blue), lifespan-extending interventions (green), and mammalian aging and mortality (red). ‘adjusted p-value < 0.1, *adjusted p-value < 0.05, **adjusted p-value < 0.01, ***adjusted p-value < 0.001. *E. Effect of ELAM treatment on predicted transcriptomic age (tAge) of mouse heart and skeletal muscle tissues.* Mouse multi-tissue transcriptomic clocks of relative chronological age (left) and mortality (right) have been applied. Benjamini-Hochberg adjusted p-values comparing young vs. old animals, and sex- and age-matched control vs. ELAM treated mice are shown in text.

Many Gene Ontology biological processing (GObp) terms were significantly impacted by ELAM treatment (Supplementary Data File 1). Therefore, to gain greater mechanistic insights, we consolidated redundant GObp terms using REVIGO^67^ and visualized the resulting interaction networks using Cytoscape^68^. For old female hearts, upregulated functions (Figure 3A) were organized into separate clusters concerning ATP generating processes, mRNA splicing, ion transport, and OXPHOS complex assembly. For old male hearts, activated functions (Figure 3B) similarly consisted of processes related to mitochondrial organization, mitochondrial transport, ATP generation, and cell communication. Despite consolidation, many different cellular functions were downregulated by ELAM treatment in both old female (Extended Data Figure 9A) and male (Extended Data Figure 9B) hearts. Again, many of these processes were related to interferon signaling and the immune system; however, additional effects on calcium/RyR2 signaling pathways were also seen.

**Figure 3:**
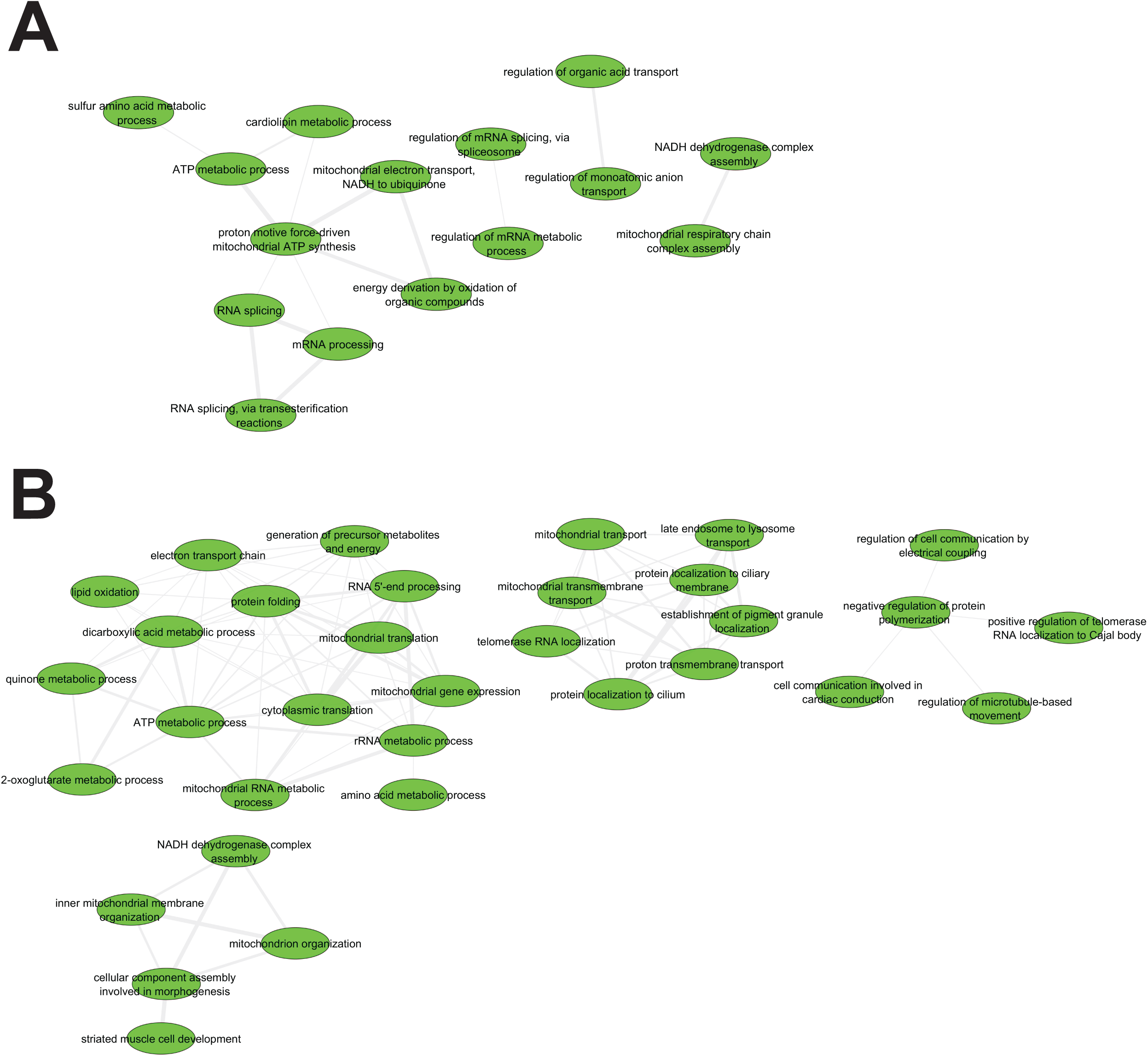
Network analysis of upregulated GObp terms following 2-months ELAM treatment in old mouse hearts. *A. Females. B. Males.* Gene Ontology biological process (GObp) terms that were significantly upregulated (FDR < 0.05) were first consolidated using REVIGO^67^ before visualization using Cytoscape^68^.

### Chronic ELAM administration does not affect tissue transcriptomic or epigenetic age

Since we observed significant attenuation of aging-related declines in muscle functional performance with ELAM, we next tested if these improvements in function could be captured by molecular biomarkers of aging. Therefore, we applied our mRNA-seq gene count data to mouse multi-tissue transcriptomic predictors of chronological age and mortality^52^ (Figure 2E). As expected, transcriptomic clocks distinguished the tissues of young and old mice (adjusted p-value < 10^−7^). Additionally, LV mass as a measure of cardiac hypertrophy was significantly positively correlated (p = 0.0028) with predicted transcriptomic age (tAge) of mouse hearts (Extended Data Figure 10A). However, only the hearts of young male mice were predicted to be biologically younger based on changes in gene expression following ELAM treatment (p = 3.5E^−5^ and p = 0.002 for the clocks of chronological age and mortality, respectively), and changes in GLS were only marginally positively correlated (p = 0.0667) with tAge (Extended Data Figure 10B). Notably, hearts of young males also showed the highest number of differentially expressed genes affected by ELAM, suggesting that the lack of tAge signal in other cases may be related to the low overall effect of ELAM on tissue transcriptomic profiles. Given that ELAM demonstrated molecular patterns associated with increased lifespan such as dynamics towards upregulated OXPHOS and mitochondrial translation, we then applied modular transcriptomic clocks to uncover if the biological age of individual cellular functions was impacted by ELAM treatment (Extended Data Figure 8B). However, none of the 23 modules had statistically significant effects on transcriptomic age under any of the treatment conditions. Thus, we determined that the impact of ELAM treatment on gene expression profiles of organs in most groups was insufficient to produce significant effects on tissue predicted biological age in a cross-sectional comparison.

To further assess whether ELAM treatment had any impact on molecular predictors of aging, we performed DNAm microarray on genomic DNA isolated from the heart tissues. By PCA (Figure 4A), we detected strong separation by sex along principal component 1, whereas the samples were not separated by either age or treatment along principal component 2. After data normalization and filtering of failed probes, we calculated mean methylation levels from the measured beta values (Figure 4B). In males, we observed a slight yet significant (p = 0.0376) increase in mean DNAm with age. However, we did not detect any impact of ELAM treatment on mean DNAm levels in either sex or age group. These results were further supported by differential methylation analysis, which revealed only 7 differentially methylated positions for young males (false discovery rate of 0.05), while no statistically significant changes in DNAm were observed for other groups and sexes. We then used epigenetic clocks to determine if ELAM had any effect on predicted biological age (Figure 4C). Using both UniversalClock2^78^ and a pan-tissue clock^79^, we measured 26-month-old male and female mice to be significantly epigenetically older than the 7-month-old mice (p < 0.0001). Moreover, we also found that the predicted tAge of the animals was well-correlated (Pearson r = 0.9151) with their respective predicted epigenetic age (Extended Data Figure 10C). However, we were unable to observe any impact of ELAM treatment on epigenetic age in either sex or in any of the other relevant clocks tested (Supplementary Data File 2). Taken together, we concluded that ELAM had virtually no impact on DNAm profiles and/or associated epigenetic age in mouse cardiac tissue.

**Figure 4:**
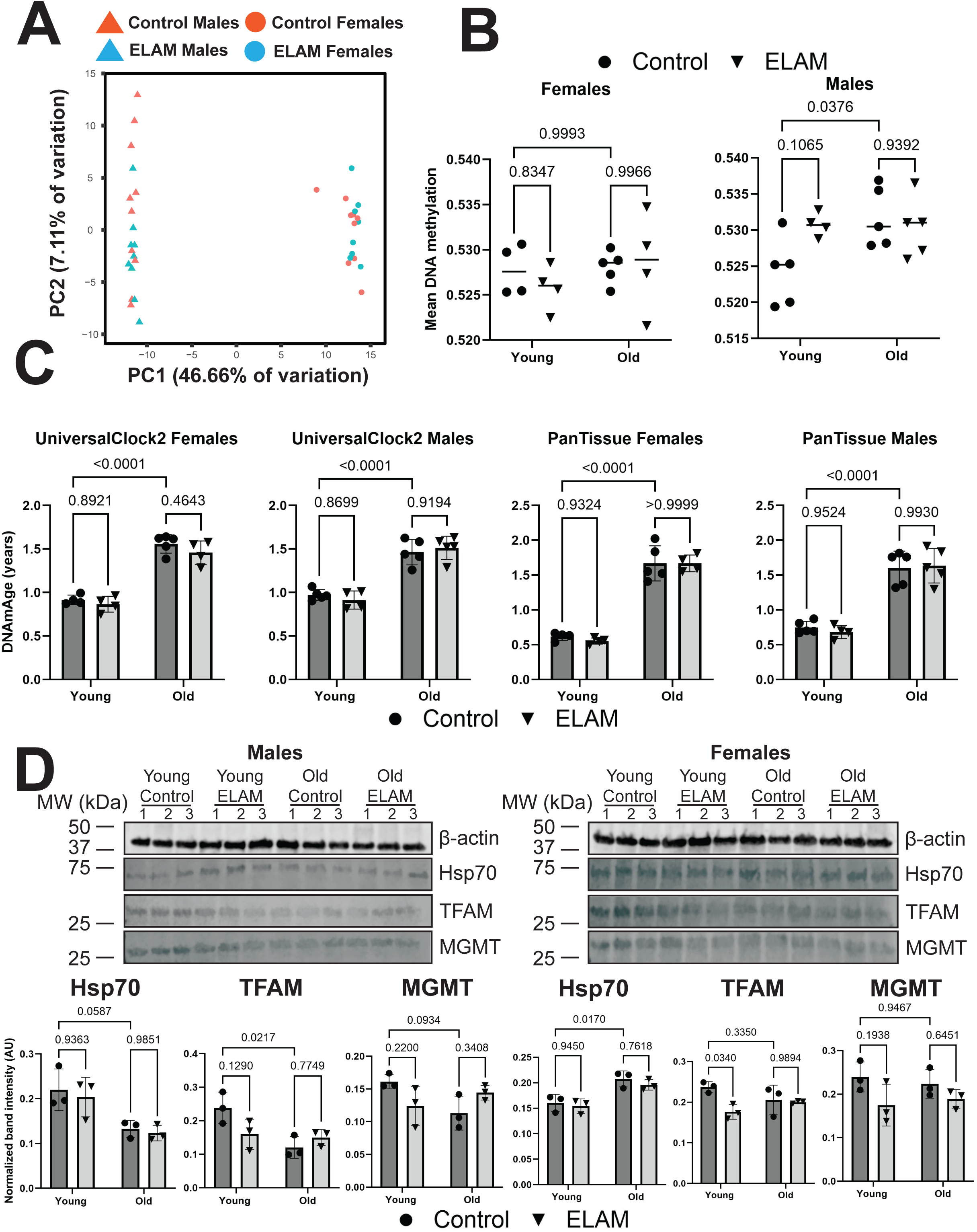
Effect of 2-months ELAM treatment on biomarkers of lifespan in young and old mouse hearts. *A. PCA of DNA methylation (DNAm) microarray samples.* PCA was performed following data normalization in SeSAMe^70^ and filtering of failed probes. *B. Mean DNAm levels.* Mean DNAm was estimated by taking the means of the filtered beta values for each sample. P-values were determined by two-way ANOVA and Tukey’s post-hoc test. P-values for the most relevant comparisons are shown in text. Lines represent sample means. *C. Effect of ELAM treatment and aging on epigenetic age (DNAmAge) of mouse hearts.* P-values were determined by two-way ANOVA and Tukey’s post-hoc test. P-values for the most relevant comparisons are shown in text. Error bars represent sample means ± standard deviations. *D. Effect of ELAM treatment and aging on protein expression of cap-independent translation (CIT) targets.* Upper panels: Western blot images of CIT targets Hsp70, TFAM, and MGMT alongside a loading control (β-actin). Lower panels: Quantification of the protein levels of CIT targets. P-values were determined by two-way ANOVA and Tukey’s post-hoc test. P-values for the most relevant comparisons are shown in text. Error bars represent sample means ± standard deviations.

Finally, to assess the potential impact of ELAM administration on alternative biomarkers of longevity, we performed Western blots for cap-independent translation (CIT) targets TFAM, MGMT, and Hsp70 in mouse hearts. Previous work done in mouse livers and kidneys has shown that the protein abundance of CIT targets decreases with age and conversely, that the upregulation of CIT targets is a shared mechanism of compounds and interventions that robustly extend mouse lifespan^80^. In males, we did observe a decrease in protein expression of CIT targets with age (Figure 4D) that was marginally significant (p = 0.0587, p = 0.0217, and p = 0.0934 for Hsp70, TFAM, and MGMT, respectively) and occurred without an effect on gene expression (Extended Data Figure 11). However, this was not observed in the hearts of female mice, which showed an increase in Hsp70 protein expression with age (p = 0.0170). Interestingly for females, TFAM was upregulated at the mRNA level in young mice following ELAM treatment (p = 0.0075) but downregulated at the protein level (p = 0.0340). TFAM is a mitochondrial transcription factor that is essential for the replication, packaging, and maintenance of mitochondrial DNA^81^ and furthermore, has been shown previously to be affected by ELAM treatment in a mouse model of Alzheimer’s disease^27^. Overall, we found that ELAM treatment appeared to have a negligible impact on several molecular predictors of mouse lifespan and longevity.

## DISCUSSION

In this study, we explored the relationship between age-related function and biological age by addressing two questions. First, we evaluated the age- and sex-specific effects of ELAM treatment on mouse frailty index, cardiac and skeletal muscle performance, and gene expression. Second, we tested if the robust improvements in skeletal and cardiac muscle function following chronic ELAM administration correlated with molecular features of aging quantified with both epigenetic and transcriptomic clocks. We found that ELAM improved general health status in longitudinal tests and mainly affected systolic function in aged males and females, whereas changes to skeletal muscle force generation were restricted to old females. At the molecular level, we observed trajectories towards upregulation of pathways related to mitochondrial organization, transport, and energy metabolism and downregulation of immune- and inflammation-related biological processes. Interestingly, despite these pathways being implicated in transcriptomic age^52^, there was no consistent effect of ELAM treatment on cardiac or skeletal muscle transcriptomic age. Moreover, in most cases, ELAM did not statistically affect the expression of any individual genes. These findings were corroborated by analyses of DNAm in mouse hearts, which showed that ELAM did not affect predicted epigenetic age or significantly alter the methylation of individual CpGs. When we limited our functional analyses to only mice whose tissues were subjected to the omics experiments, we still observed statistically significant effects on post-treatment frailty and GLS in old female and male mice, respectively. Thus, our study has demonstrated that improvements in age-related biological and tissue functions following a treatment may occur without measurable changes in biological age, as quantified with validated transcriptomic and epigenetic clocks at the study endpoint. More generally, these findings indicate that age-related function can be partially uncoupled from molecular biological age. Even though they are often corroborated in aging studies, changes in age-related function do not necessarily equate to changes in the underlying molecular drivers of aging. Furthermore, directly targeting a “secondary” or downstream hallmark of aging^82^ may not have the same impact on biological age as targeting upstream growth or developmental-related pathways.

ELAM is suggested to localize within mitochondria and increase OXPHOS efficiency and the coupling of membrane potential generation with ATP synthesis as its central mechanism of action^83^. Two non-mutually exclusive and partially overlapping explanations for this mechanism have been proposed. In the first, ELAM is targeted to mitochondria due to preferential binding interactions with their cardiolipin (CL)-rich membranes, which feature a strong negative surface charge^23,39^. As a polybasic peptide, binding of ELAM to CL partially attenuates the membrane surface potential; moreover, membrane insertion of its hydrophobic sidechains alters lipid packing interactions and membrane exposure to water. Together, these interactions may reduce excessive mitochondrial calcium signaling (a known instigator of mitochondrial dysfunction^84,85^) by displacing calcium from the membrane surface and modulate membrane curvature, which affects OXPHOS complex assembly and mitochondrial bioenergetic function^86,87^. In the second, direct binding interactions of ELAM with proteins involved in mitochondrial energy generation increases mitochondrial ATP generating capacity and coupling efficiency^36,61^. In particular, proton leak through the ANT protein has been identified as a potential cause of aging-related cardiac mitochondrial dysfunction, and treatment with ELAM is able to reduce it^37,38^.

Mitochondrial dysfunction is an established contributor to sarcopenia and heart failure^88,89^. During heart failure in humans, energy metabolism switches from primarily fatty acid beta-oxidation to include more utilization of glucose^90^. This coincides with disrupted OXPHOS and an inability to meet ATP energy requirements^91^. In turn, this lack of ATP contributes to alterations in calcium signaling, which can overload mitochondria with calcium via phosphorylation of RyR2^92^. Finally, this increase in mitochondrial calcium levels can trigger opening of the mitochondrial permeability transition pore, which further depletes ATP levels, increases mitochondrial swelling and reactive oxygen species production, and can lead to ischemia-reperfusion injury and cell death^93,94^. Therefore, ELAM treatment may improve aging-related declines in cardiac function by increasing inner membrane curvature, improving OXPHOS efficiency, and increasing cellular ATP production. Alternatively, ELAM may have direct binding interactions with ATP synthase dimers, which have been suggested to constitute the pore-forming component of the mitochondrial permeability transition pore^36,95^. Thus, ELAM binding to ATP synthase dimers might decrease their sensitivity to calcium-induced pore formation^23,24,37^.

Biomarkers of aging, such as epigenetic, transcriptomic, and proteomic aging clocks, are emerging tools to quantify biological age, assess the impacts of disease states, and perform large-scale unbiased screens to identify interventions that affect aging. However, the interpretability of their predictions is often complicated^53–56^. Recent studies have partially resolved this issue by separating adaptive CpG sites from damaging sites^48^, and by breaking down conventional clocks into specific biological modules associated with various cellular functions^52^. At the same time, it still remains unclear how biological age correlates with various tissue functions and parameters of organismal physiological health. If measurements of functional changes disagree with their predicted biological ages, how should this be reconciled? Moreover, if interventions are tested for their impact on both physiological function and molecular features of aging, and that these assessments disagree with one another, should aging be viewed as affected by these interventions? In the case of our study, we observed improvements in age-related function following ELAM treatment that were not consistently detected by epigenetic or transcriptomic predictors of biological age. Not only are healthspan and lifespan two separate parameters^96^, but they also may be indirectly related to changes in function, and all three may be different from changes in biological age. In support of this, albeit with a limited sample size and different route of administration, ELAM treatment was previously shown not to affect mouse lifespan^42^. However, we also cannot rule out the possibility that molecular clocks based on individual modalities may not represent certain changes induced at other levels of tissue organization (e.g., concentration of metabolites and glycans)^97^, that the effect of ELAM on heart epigenetic and transcriptomic age may be masked by the fact that cardiomyocytes only make up around 25-35% of the total number of cells^98^, or that functional changes respond more quickly to improved mitochondrial function and a longer treatment may be required to remodel the underlying molecular markers of biological age.

In agreement with previous studies^31,38,61^, we have shown that ELAM treatment can mitigate some age-related declines in mouse physiological muscle function. Improvements in function were observed in both sexes, which is important considering that most ITP drugs are more effective in males than females^6,7,10^. Furthermore, the proposed mechanisms of ELAM are distinct from those of known geroprotectors, which traditionally affect developmental and/or growth-related pathways to alter aging trajectories regardless of the age at which treatment is initiated. ELAM seems to be unique as it is proposed to affect only cells that have unhealthy and/or dysfunctional mitochondria^27,99,100^, including cells from aged tissues^37^. Finally, as a peptide drug candidate, ELAM is safe and well-tolerated but traditionally requires SQ administration. Taken together, our study confirms ELAM as a potential novel intervention for mitigating aging-related declines in tissue function.

### Future Directions and Limitations

In future studies, it would be interesting to compare the effects of other mitochondria-targeted treatments on tissue function and predicted biological age to see if downstream pathways such as mitochondrial metabolism have less effect on biological age compared to more universal upstream pathways related to growth and development. Additionally, it would be important to test other peptide analogs to see if they can more strongly attenuate aging-related declines in cardiac and skeletal muscle performance. In the current study, we utilized inbred C57BL/6J mice; as such, further validation may be necessary to determine if the effects of chronic ELAM on muscle function are consistent in other strains and genetically diverse populations. The sample sizes for the omics experiments are limited to n = 4-5 per treatment condition. Finally, additional aging biomarkers may be applied to assess biological functions of cells and tissues in response to ELAM and related interventions.

## ACKNOWLEDGEMENTS

ELAM was provided by Stealth BioTherapeutics Inc. at no cost through an existing material transfer agreement. Stealth BioTherapeutics Inc. had no role in the study design or analysis of the data. The authors would like to thank Alice Kane for training and assistance with experimental methods.

## AUTHOR CONTRIBUTION

Created study design: W.M, G.P, D.J.M, V.N.G

Conducted experiments: W.M, G.P

Analyzed experimental data: W.M, G.P, A.T

Oversaw experiments and provided laboratory resources: D.J.M, V.N.G

Wrote the initial draft: W.M, G.P

Revised the manuscript: W.M, G.P, A.T, M.C, D.J.M, V.N.G

---

**Extended Data Figure 1: Similar components of the 31-point frailty index are elevated in old male and female mice.** Heatmap of average value of each component of the frailty index in 4-5-month-old (young) female and male (n=9-10) and 23-24-month-old (old) female (n=23-24) at baseline. Only mice that survived to study endpoint were included in the analysis.

**Extended Data Figure 2: Aging and elamipretide treatment analyzed by echocardiography strain analysis.** *Baseline echocardiography strain analysis of parasternal long axis images of the LV of young (Yng) and old mice for A. End diastolic volume (EDV), B. End systolic volume (ESV), C. Stroke Volume (SV), D. Fractional shortening (FS), E. Cardiac output (CO), F. End diastolic left ventricular mass (EDLVM), G. End systolic left ventricular mass (ESLVM), and H. Heart rate. Δ (Post-pre) echocardiography strain analysis of parasternal long axis images of the LV of aged control (Con) and ELAM treated mice for I. EDV, J. ESV, K. SV, L. FS, M. CO, N. EDLVM, O. ESLV), and P. Heart rate.* 4-5-month-old (young) female and male (n=9-10) and 23-24-month-old (old) female (n=23-24) and male were compared for each measurement at baseline. ELAM treatment effects were compared in control and ELAM-treated old mice (n=11-12) using Δ measurements. Only mice that survived to study endpoint were included in the analysis. Statistical significance was determined by two-way ANOVA with Tukey’s post-hoc test. Significant ANOVA factors written in text with selected Tukey’s post hoc test comparisons on graphs. Error bars represent sample means ± standard deviations.

**Extended Data Figure 3: Systolic function in aging and elamipretide treatment analyzed by conventional echocardiography.** *Baseline conventional echocardiography systolic function quantified from the short axis M-mode images of the LV of young (Yng) and old mice for A. End systolic diameter (ESD), B. End diastolic diameter (EDD), C. ESV, D. EDV, E. SV, F. Ejection fraction (EF), G. FS, and H. CO. Δ (Post-pre) conventional echocardiography systolic function quantified from the short axis M-mode images of the LV of aged control (Con) and ELAM treated mice for I. ESV, J. EDD, K. ESV, L. EDV, M. SV, N. EF, O. FS, and P. CO.* 4-5-month-old (young) female and male (n=9-10) and 23-24-month-old (old) female (n=23-24) and male were compared for each measurement at baseline. ELAM treatment effects were compared in control and ELAM-treated old mice (n=11-12) using Δ measurements. Only mice that survived to study endpoint were included in the analysis. Statistical significance was determined by two-way ANOVA with Tukey’s post-hoc test. Significant ANOVA factors written in text with selected Tukey’s post hoc test comparisons on graphs. Error bars represent sample means ± standard deviations.

**Extended Data Figure 4: Diastolic function in aging and elamipretide treatment analyzed by conventional echocardiography.** *Baseline conventional echocardiography diastolic function quantified from 4-chamber tissue doppler and pulse wave doppler images of the LV of young (Yng) and old mice for A. Mitral valve (MV) E wave, B. MV A wave, C. E’ wave, D. A’ wave, E. MV E/E’, F. MV E/A, G. E’/A’, and H. A’/E’. Δ (Post-pre) conventional echocardiography diastolic function quantified from 4-chamber tissue doppler and pulse wave doppler images of the LV of aged control (Con) and ELAM treated mice for I. MV E wave, J. MV A wave, K. E’ wave, L. A’ wave, M. MV E/E’, N. MV E/A, O. E/A’, P. A’/E’.* 4-5-month-old (young) female and male (n=9-10) and 23-24-month-old (old) female (n=23-24) and male were compared for each measurement at baseline. ELAM treatment effects were compared in control and ELAM-treated old mice (n=11-12) using Δ measurements. Only mice that survived to study endpoint were included in the analysis. Statistical significance was determined by two-way ANOVA with Tukey’s post-hoc test. Significant ANOVA factors written in text with selected Tukey’s post hoc test comparisons on graphs. Error bars represent sample means ± standard deviations.

**Extended Data Figure 5: Cardiac hypertrophy in aging and elamipretide treatment analyzed by conventional echocardiography.** *Baseline conventional echocardiography cardiac hypertrophy from the short axis M-mode images of the LV of young (Yng) and old mice for A. LV Mass, B. LV Mass Cor (corrected), C. Left ventricular anterior wall thickness in systole (LVAW;s), D. Left ventricular anterior wall thickness in diastole (LVAW;d), E. Left ventricular posterior wall thickness in systole (LVPW;s), F. Left ventricular posterior wall thickness in diastole (LVPW;d). At study endpoint, the heart was collected and weighed for control young and old mice for G. Heart mass, and H. Heart mass normalized to tibia length. Δ (Post-pre) conventional echocardiography cardiac hypertrophy from the short axis M-mode images of the LV of aged control (Con) and ELAM treated mice for I. LV Mass, J. LV Mass Cor, K. LVAW;s, L. LVAW;d, M. LVPW;s, N. LVPW;d.. At study endpoint, the heart was collected and weighed for control and ELAM treated old mice for G. Heart mass, and H. Heart mass normalized to tibia length.* 5-month-old (young) female and male (n=9-10) and 23-24-month-old (old) female (n=23-24) and male were compared for each measurement at baseline. ELAM treatment effects were compared in control and ELAM-treated old mice (n=11-12) using Δ measurements. Only mice that survived to study endpoint were included in the analysis. Heart masses were collected at study endpoint from 7 month (young) and 26 month (old) male and female control and ELAM treated mice. Statistical significance was determined by two-way ANOVA with Tukey’s post-hoc test. Significant ANOVA factors written in text with selected Tukey’s post hoc test comparisons on graphs. Error bars represent sample means ± standard deviations.

**Extended Data Figure 6: *In vivo* muscle force-frequency in aging and elamipretide treatment.** *At study endpoint, the gastrocnemius muscles were collected, weighed, and normalized to tibia length for A. Control young and old mice, and B. Control and ELAM treated old mice. C. Baseline in vivo muscle force normalized to body mass across a range of stimulation frequencies in young and old mice. D. Baseline maximum in vivo muscle force normalized to body mass of young and old mice. E. Δ (Post-pre) in vivo muscle force normalized to body mass across a range of stimulation frequencies at endpoint for old mice. F. Δ maximum in vivo muscle force normalized to body mass at endpoint for old mice. G. Baseline maximum rate of muscle contraction across a range of stimulation frequencies in young and old mice. H. Δ maximum rate of muscle contraction across a range of stimulation frequencies at endpoint for old control and ELAM mice. I. Baseline maximum rate of muscle relaxation across a range of stimulation frequencies in young and old mice. J. Δ maximum rate of muscle relaxation across a range of stimulation frequencies at endpoint for old control and ELAM treated mice.* 5-month-old (young) female and male (n=9-10) and 23-24-month-old (old) female (n=23-24) and male were compared for each measurement at baseline. ELAM treatment effects were compared in control and ELAM-treated old mice (n=11-12) using Δ measurements. Only mice that survived to study endpoint were included in the analysis. Gastrocnemius muscle masses were collected at study endpoint from 7 month (young) and 26 month (old) male and female control and ELAM treated mice. Statistical significance was determined by two-way ANOVA with Tukey’s post-hoc test except for comparisons that include stimulation frequency as a factor, which were analyzed by three-way ANOVA. Significant ANOVA factors written in text with selected Tukey’s post hoc test comparisons on graphs. Error bars represent sample means ± standard deviations.

**Extended Data Figure 7: *In vivo* muscle fatigue contraction and relaxation in aging and elamipretide treatment.** *A. Baseline maximum rate of muscle contraction across 120 fatiguing stimulations in young and old mice. B. Δ (Post-pre) maximum rate of muscle contraction across 120 fatiguing stimulations at endpoint for old control and ELAM treated mice. C. Baseline maximum rate of muscle relaxation contraction across 120 fatiguing stimulations in young and old mice. D. Δ maximum rate of muscle relaxation contraction across 120 fatiguing stimulations at endpoint for old control and ELAM treated mice.* 5-month-old (young) female and male (n=9-10) and 23-24-month-old (old) female (n=23-24) and male were compared for each measurement at baseline. ELAM treatment effects were compared in control and ELAM-treated old mice (n=11-12) using Δ measurements. Only mice that survived to study endpoint were included in the analysis. Statistical significance was determined by three-way ANOVA with Tukey’s post-hoc test. Significant ANOVA factors written in text with selected Tukey’s post hoc test comparisons on graphs. Error bars omitted for clarity.

**Extended Data Figure 8: Effect of 2-months ELAM treatment on aging-related changes in gene expression.** *A. Correlation analysis of gene expression changes induced by ELAM treatment at the level of individual genes (left) and enriched pathways (right).* Spearman correlation between normalized enrichment scores (NES) from gene set enrichment analysis (GSEA) performed for signatures of ELAM treatment (blue), lifespan-extending interventions (green), and mammalian aging and mortality (red). ‘adjusted p-value < 0.1, *adjusted p-value < 0.05, **adjusted p-value < 0.01, ***adjusted p-value < 0.001. *B. Modular transcriptomic clock analysis.* Normalized difference in tAge between control and ELAM-treated mice estimated with individual module-specific transcriptomic clocks of chronological age and mortality. Negative (blue) and positive (red) values represent reduced and elevated tAge in ELAM-treated animals, respectively. No statistically significant differences (Benjamini-Hochberg adjusted p-value < 0.05) were observed for any modular clocks.

**Extended Data Figure 9: Network analysis of downregulated GObp terms following 2-months ELAM treatment in old mouse hearts.** *A. Females. B. Males.* Gene Ontology biological process (GObp) terms that were significantly upregulated (False Discovery Rate < 0.05) were first consolidated using REVIGO^67^ before visualization using Cytoscape^68^.

**Extended Data Figure 10: Pearson correlation analyses.** *A. Correlation of LV mass with tAge.* Data points correspond to LV mass (post-treatment) and tAge (relative) of hearts from young and old male and female mice (both treatments). *B. Correlation of tAge with global longitudinal strain (GLS).* Data points correspond to ΔGLS (Post-pre) and tAge (relative) of hearts from old male and female mice (both treatments). *C. Correlation of DNAmAge with tAge.* Data points correspond to DNAmAge (PanTissue) and tAge (relative) of hearts from young and old male and female mice (both treatments).

**Extended Data Figure 11: Normalized gene expression of CIT targets.** *A. Hsp70. B. TFAM. C. MGMT.* Benjamini-Hochberg adjusted p-values comparing sex- and age-matched control vs. ELAM treated mice are shown in text.

