## Extended Data Figures for "The mitochondrial-targeted peptide therapeutic elamipretide improves cardiac and skeletal muscle function during aging without detectable changes in tissue epigenetic or transcriptomic age"

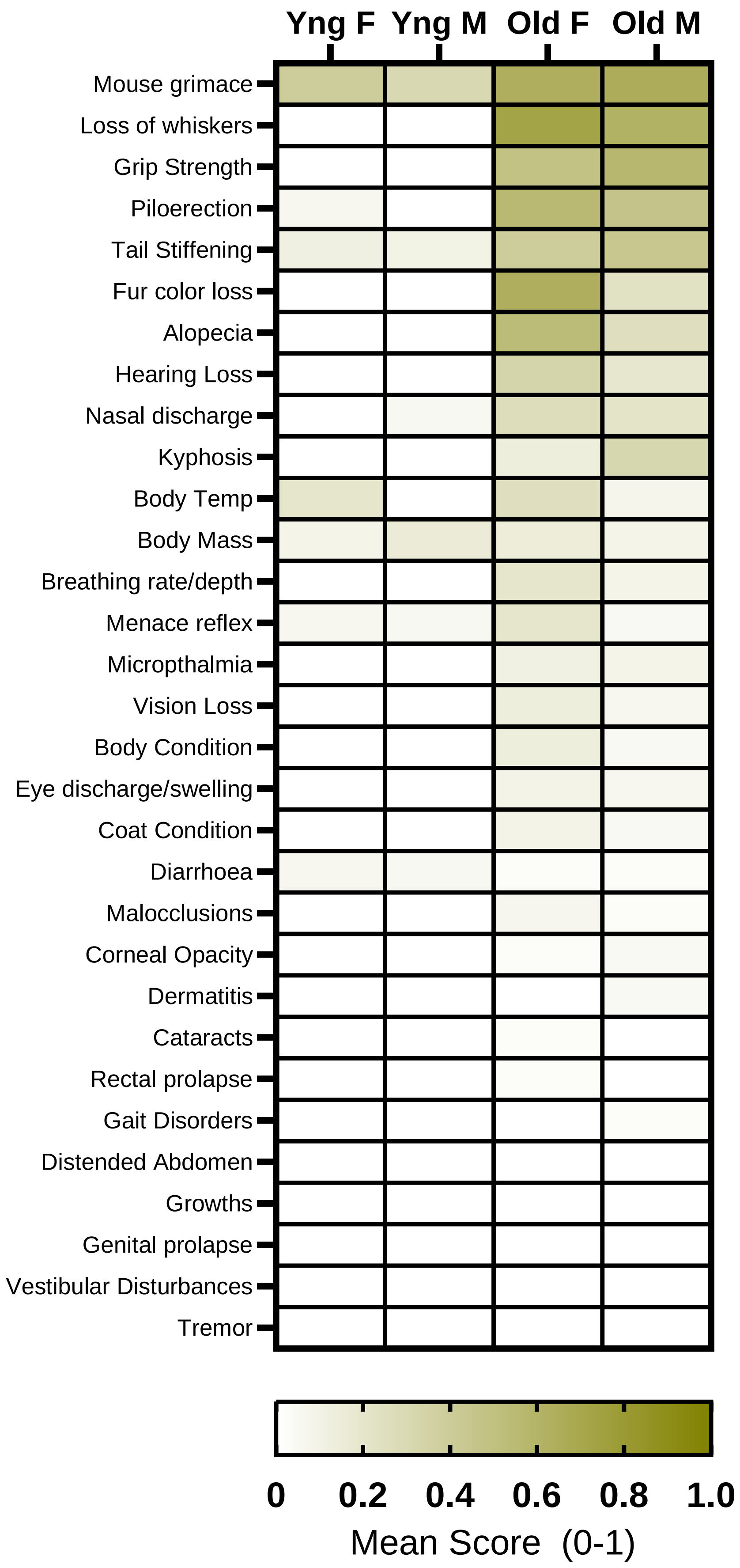


### Extended Data Figure 1

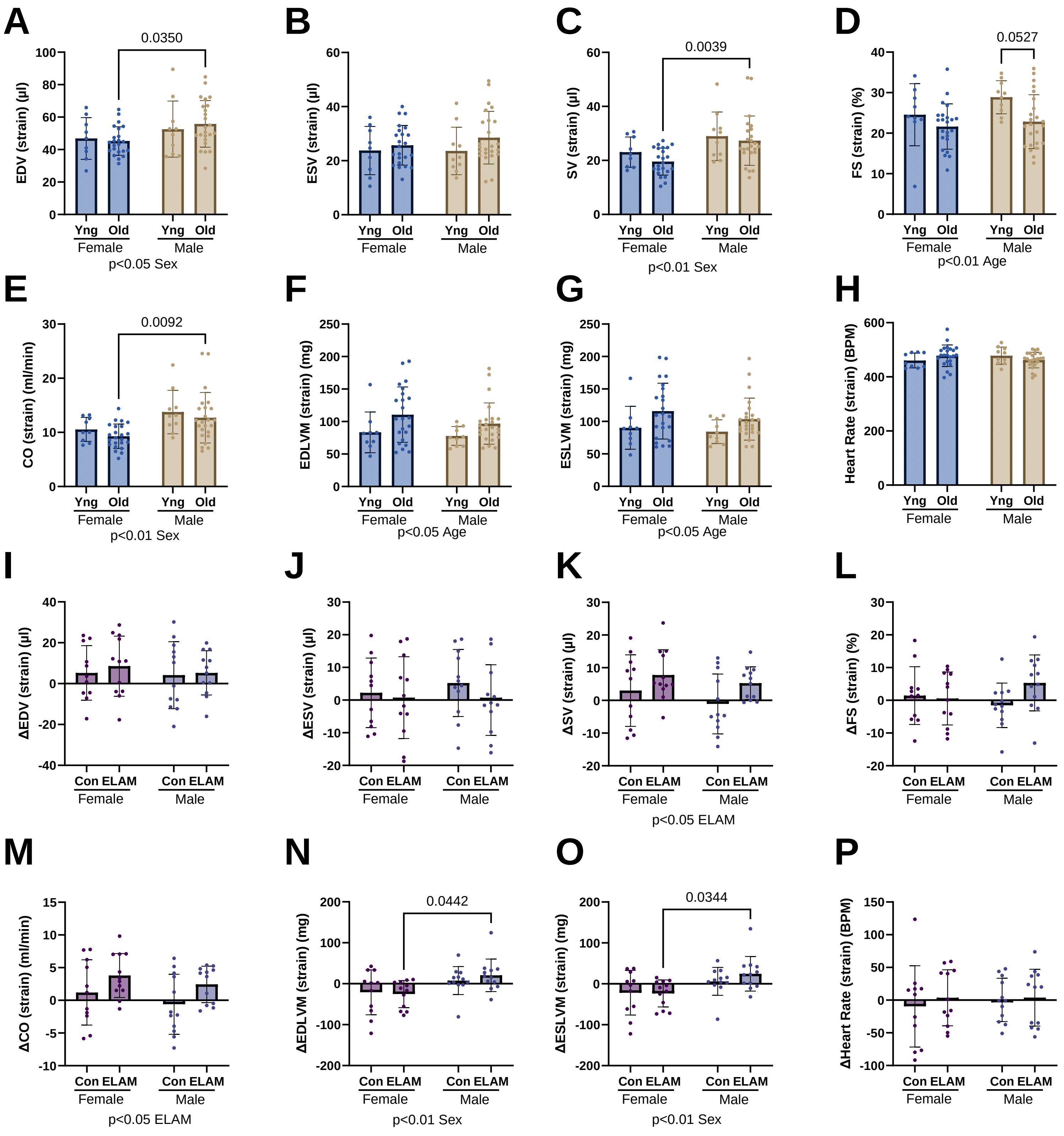


### Extended Data Figure 2

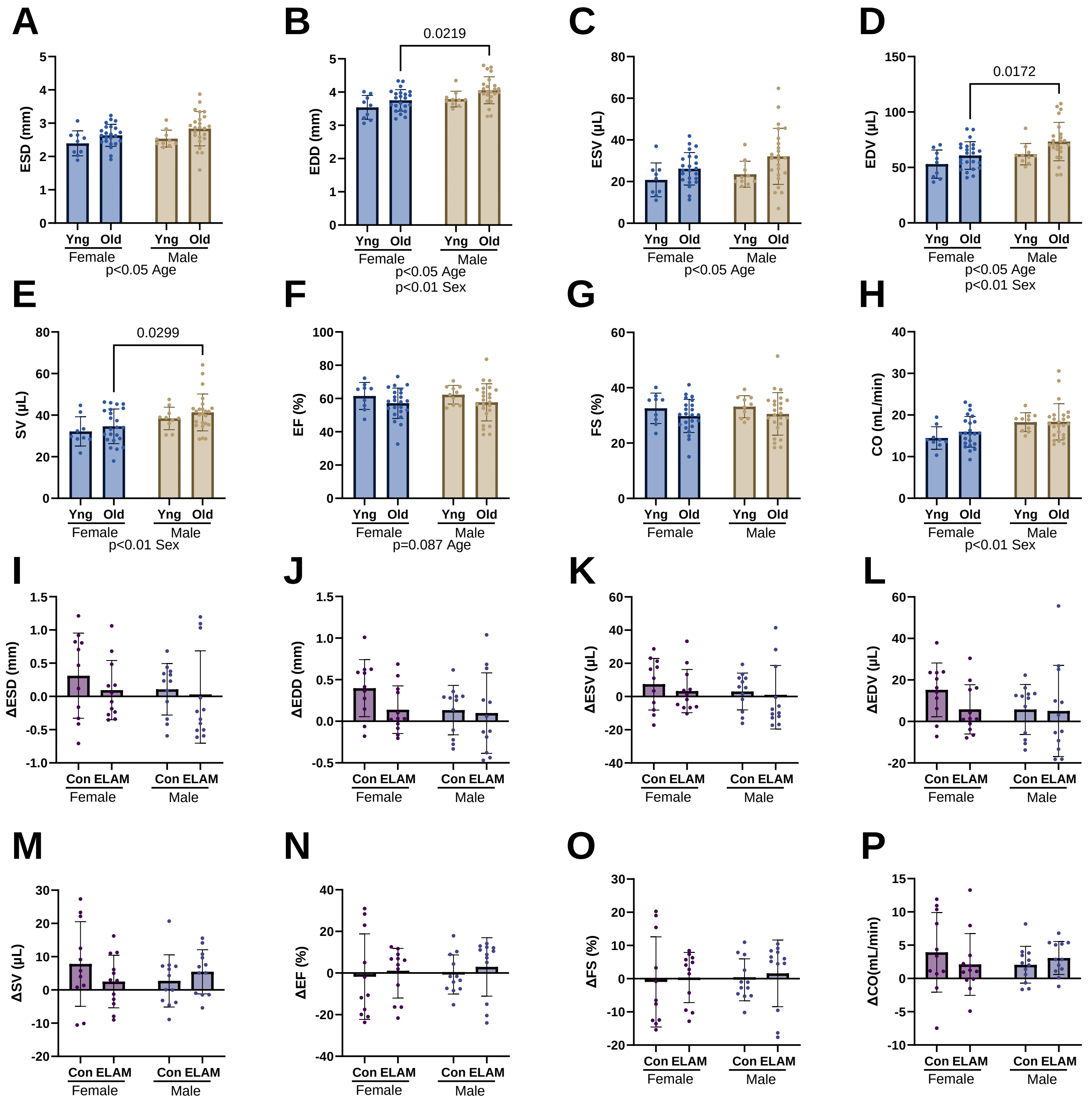


### Extended Data Figure 3

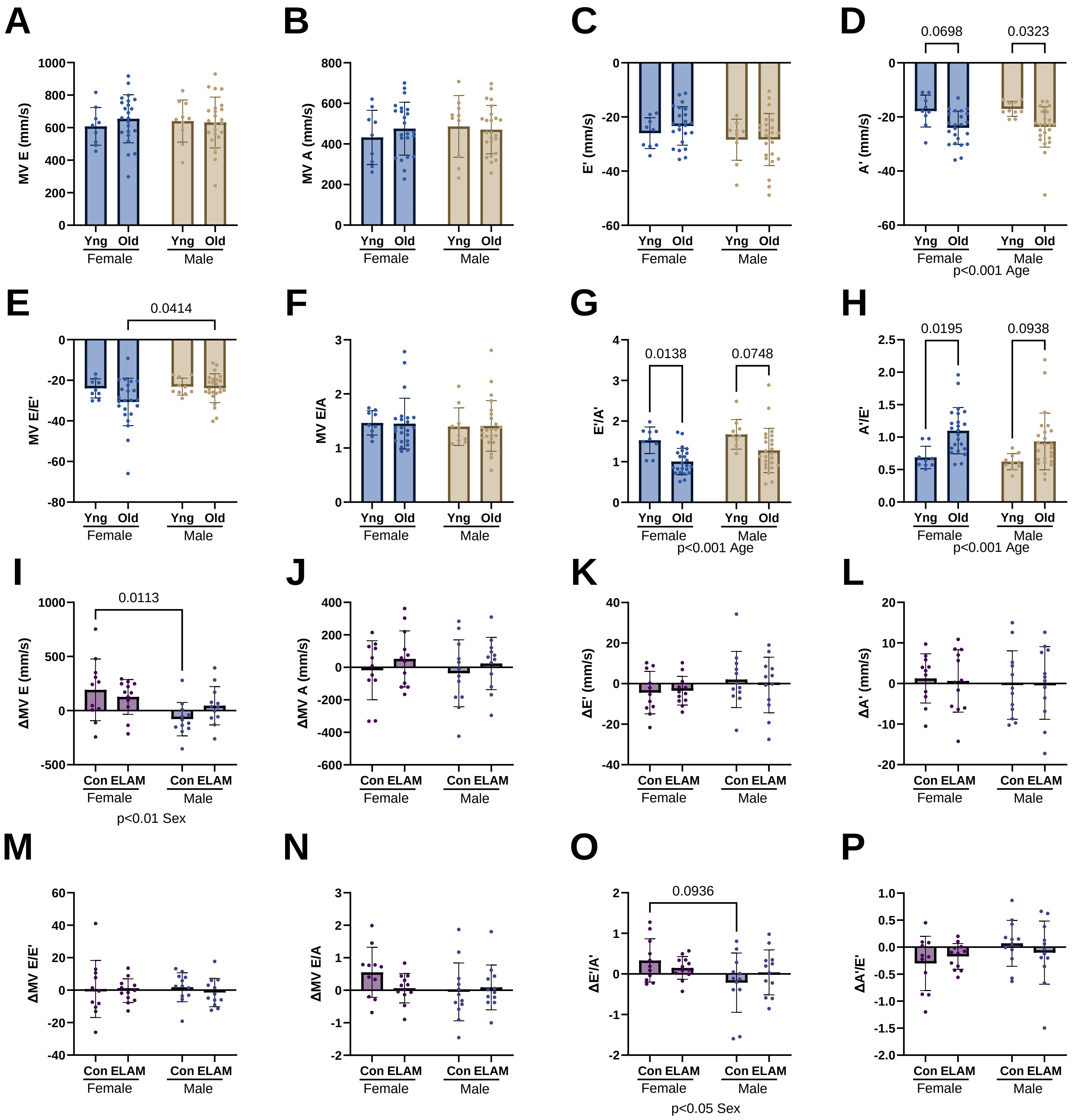


### Extended Data Figure 4

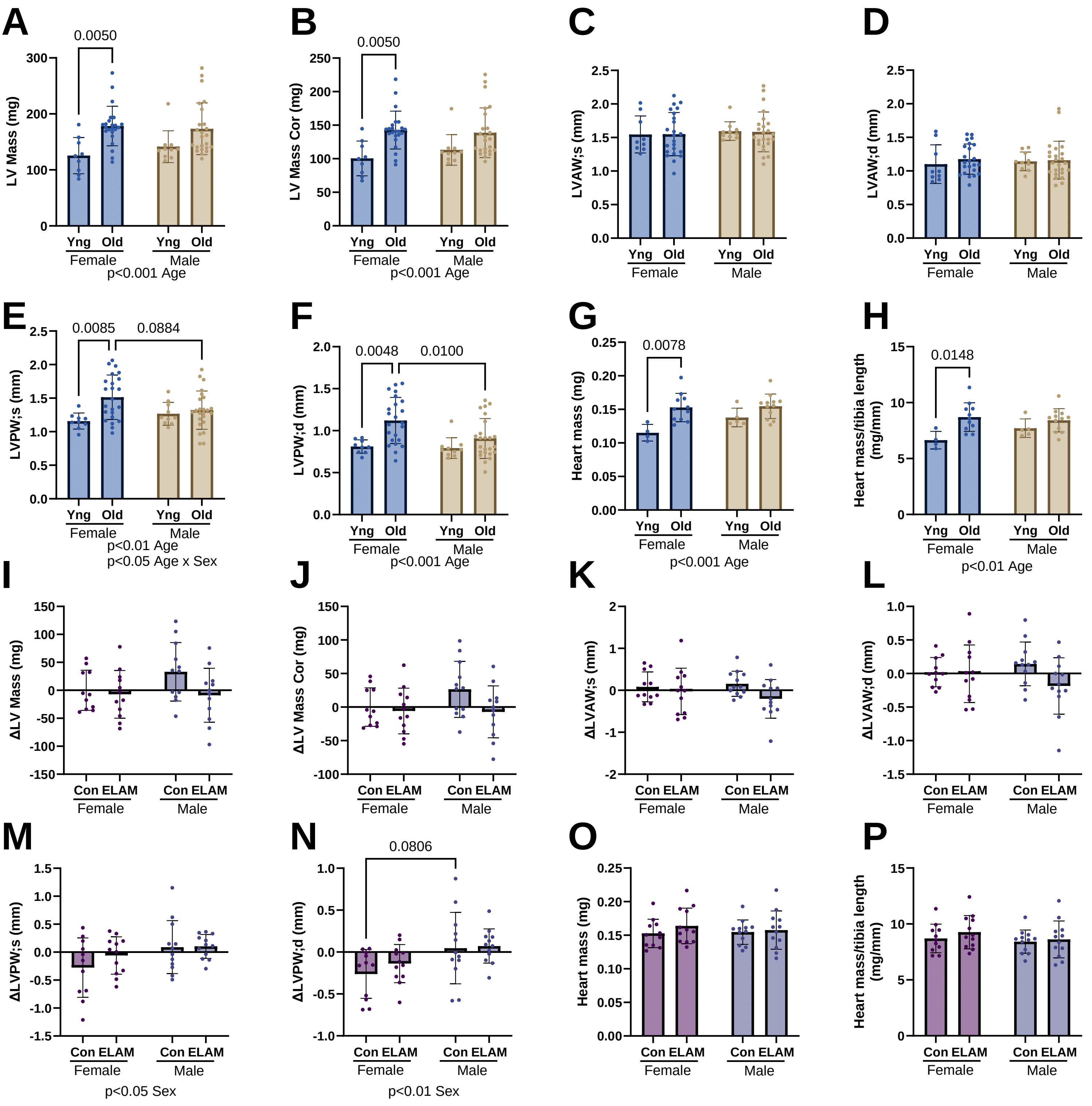


### Extended Data Figure 5

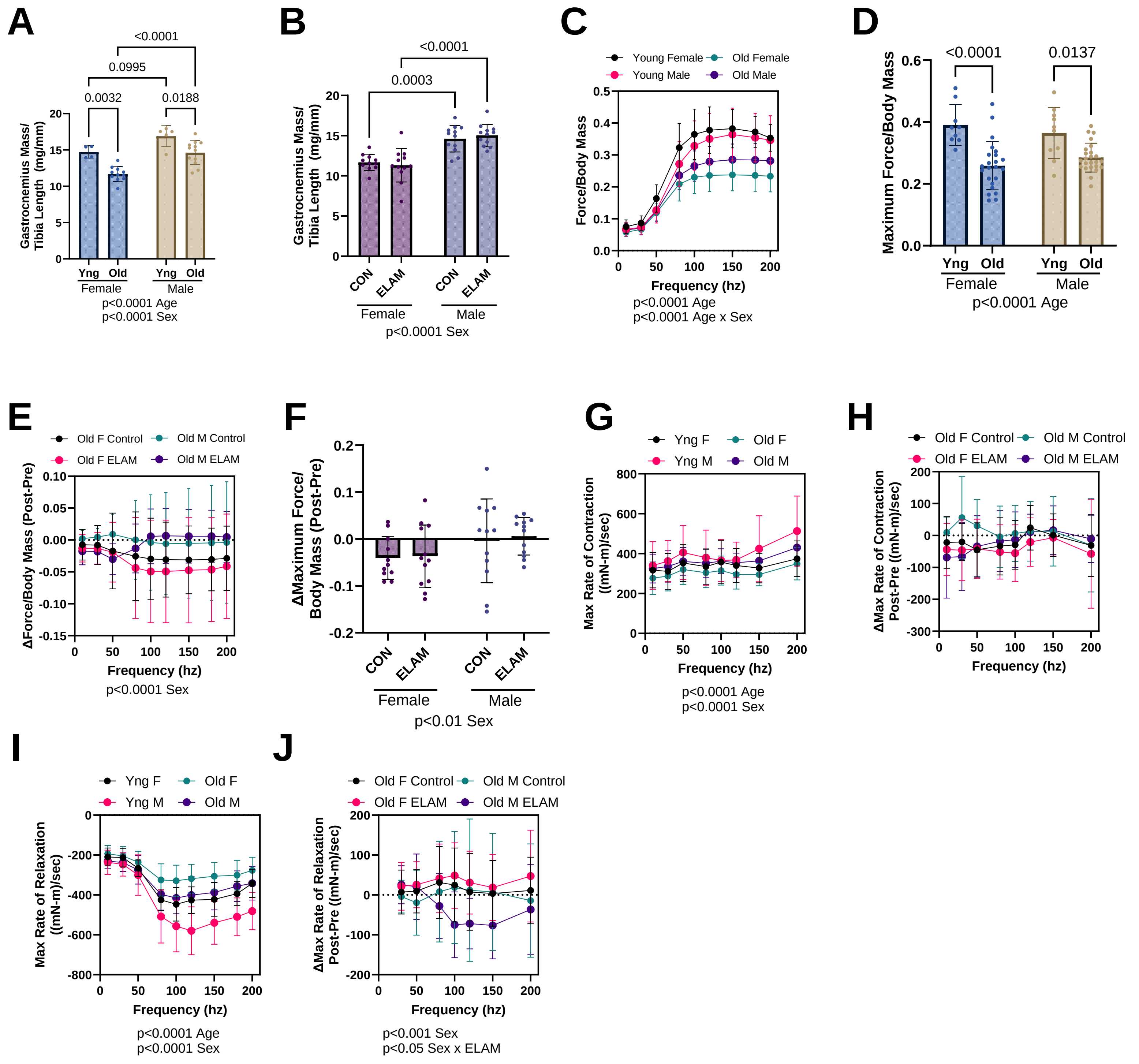


### Extended Data Figure 6

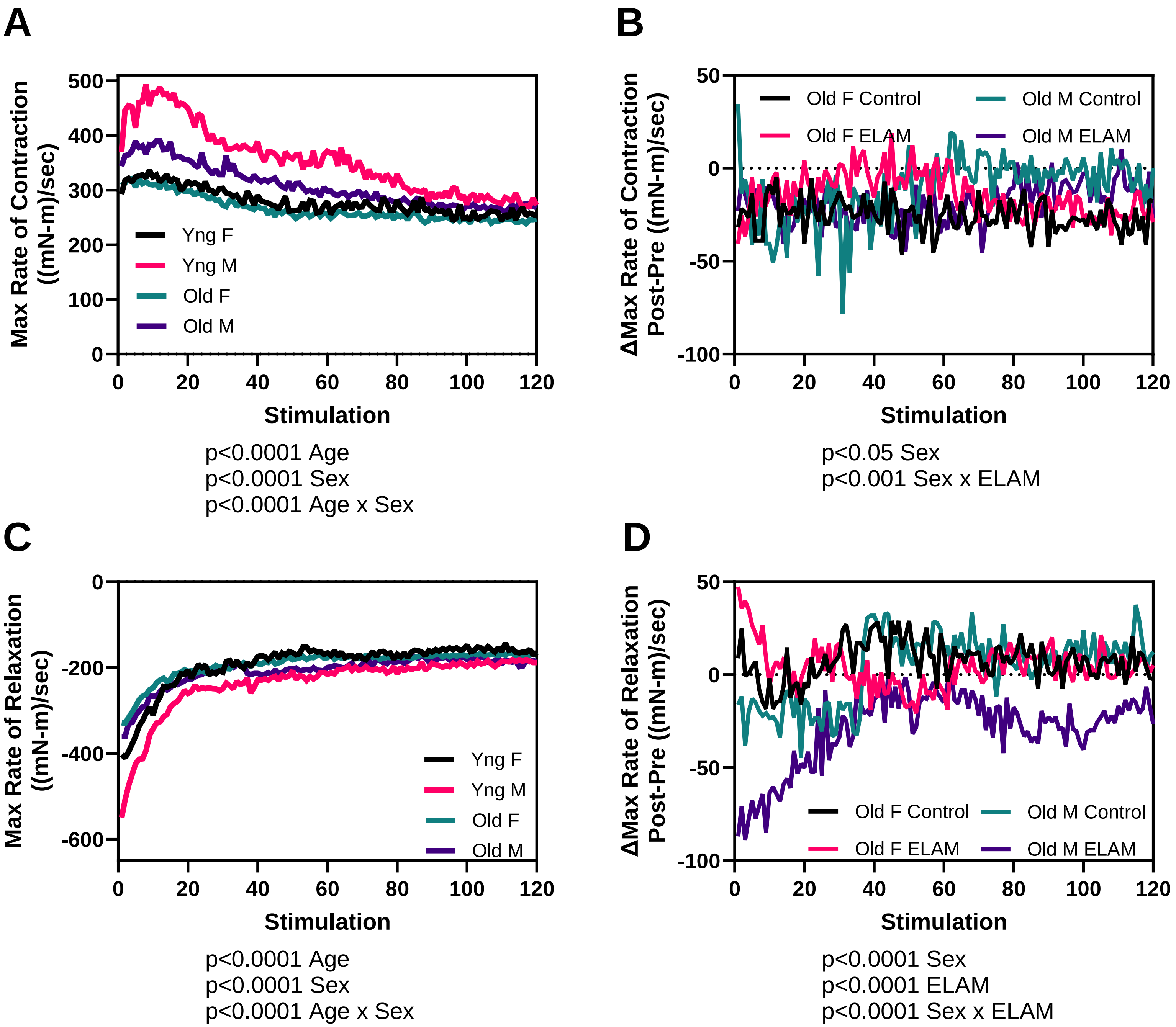


### Extended Data Figure 7

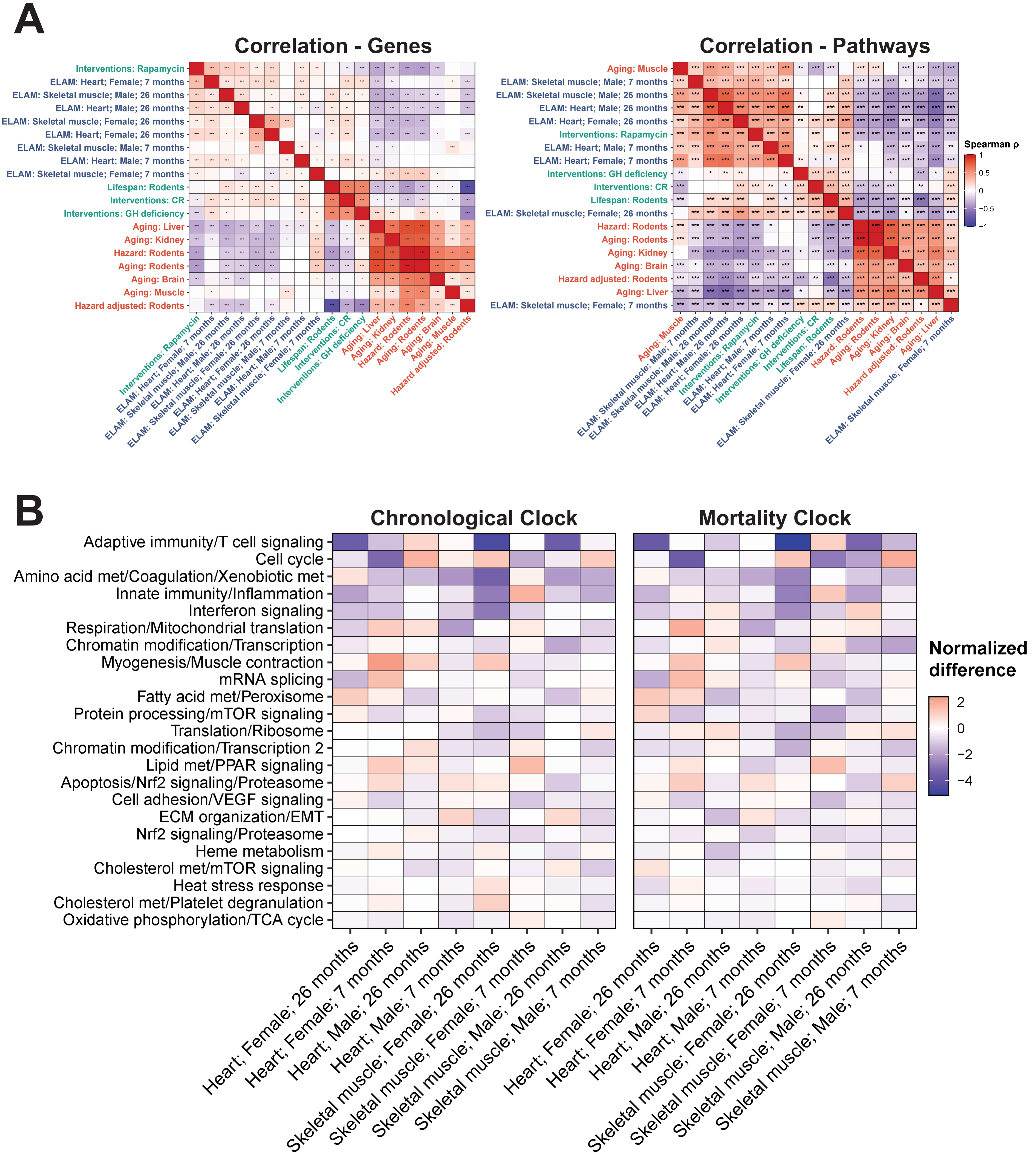


### Extended Data Figure 8

# A

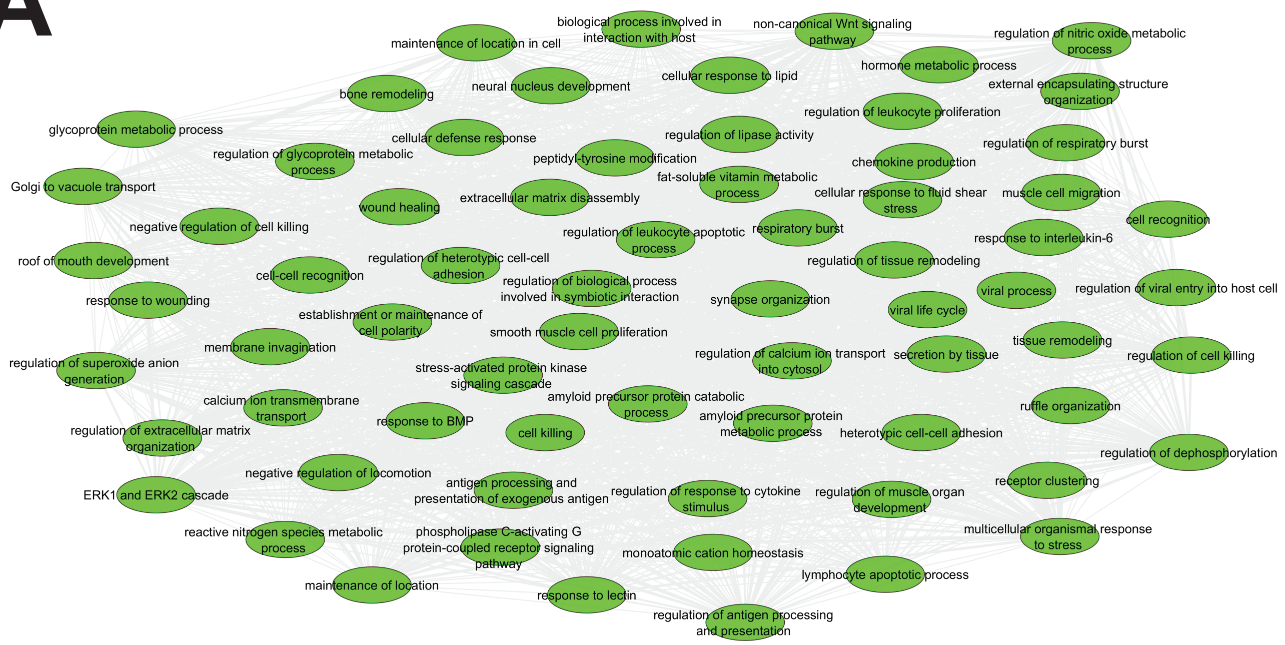


# B

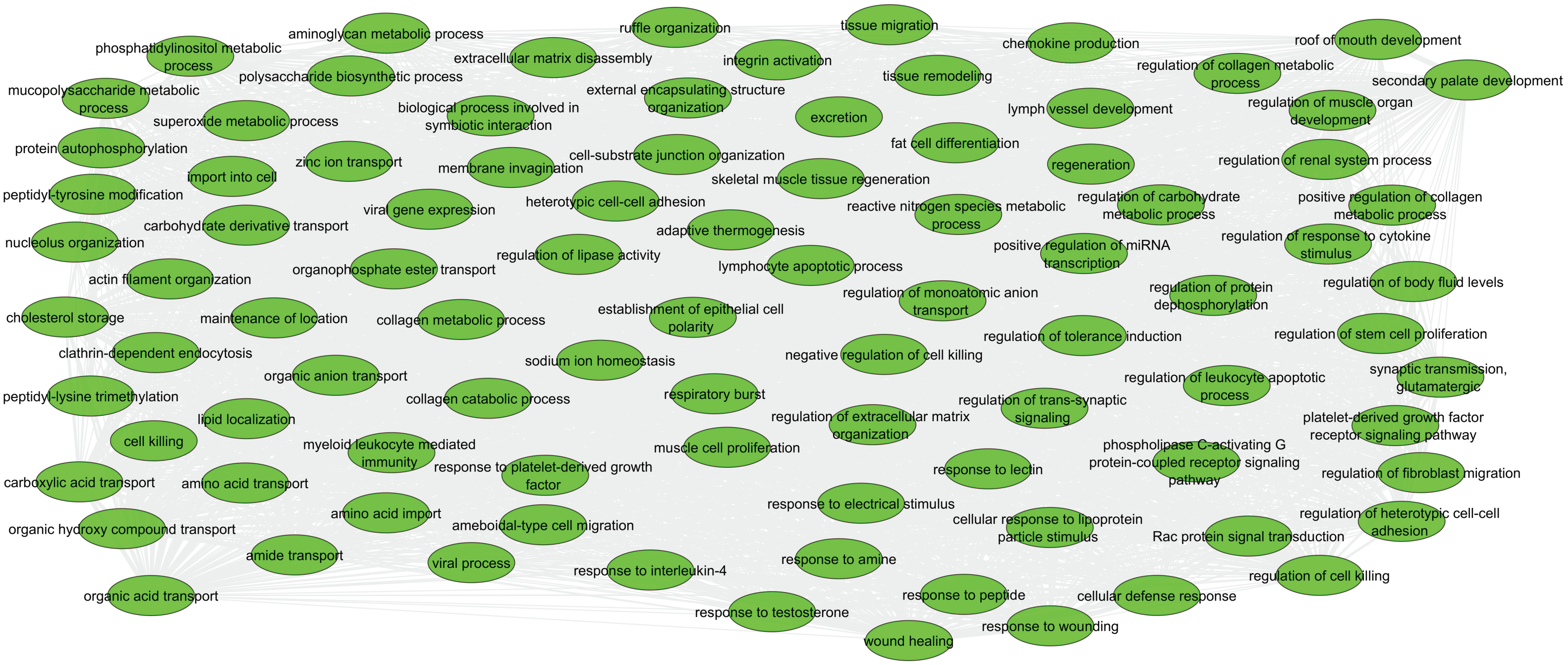


### Extended Data Figure 9

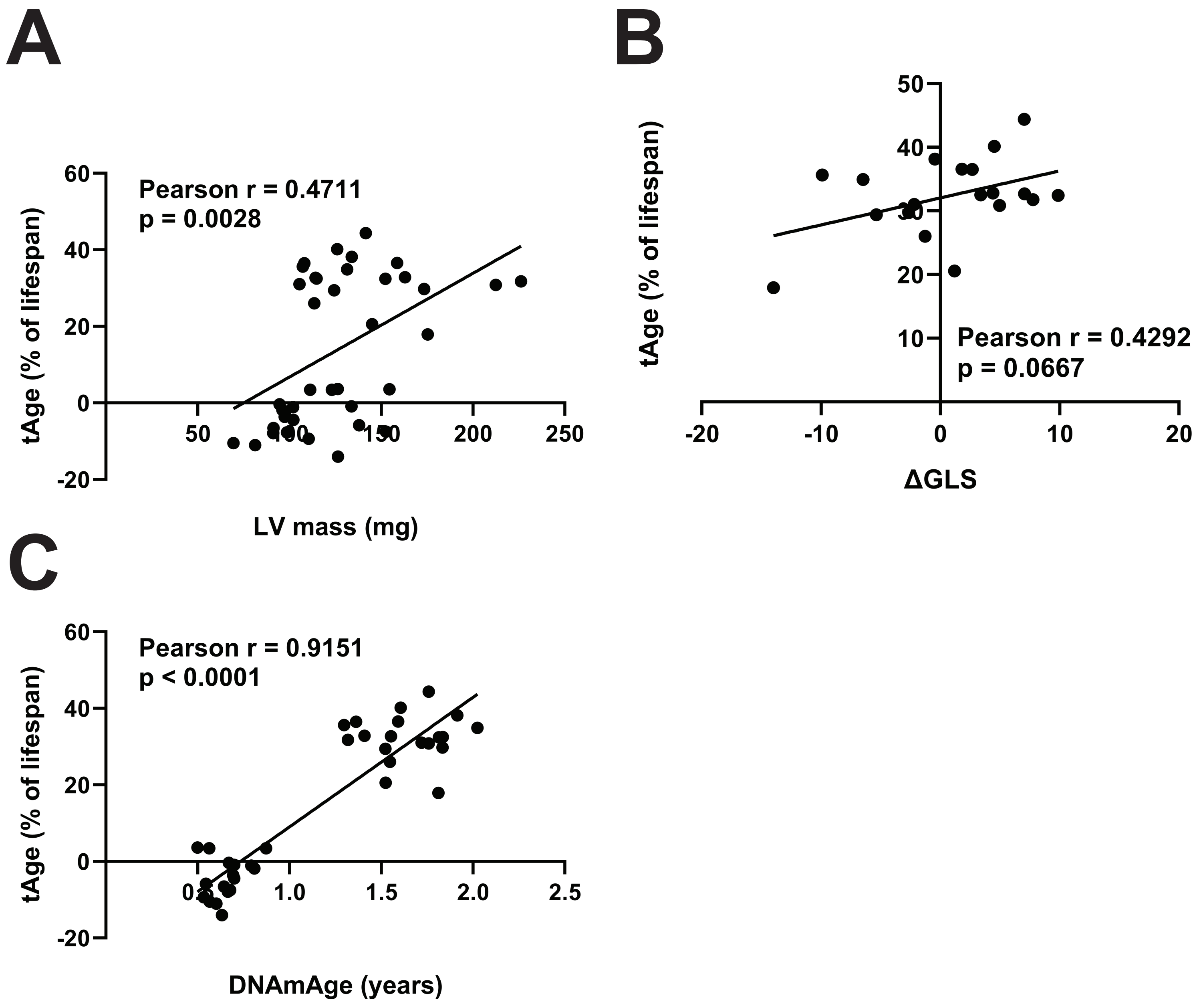


### Extended Data Figure 10

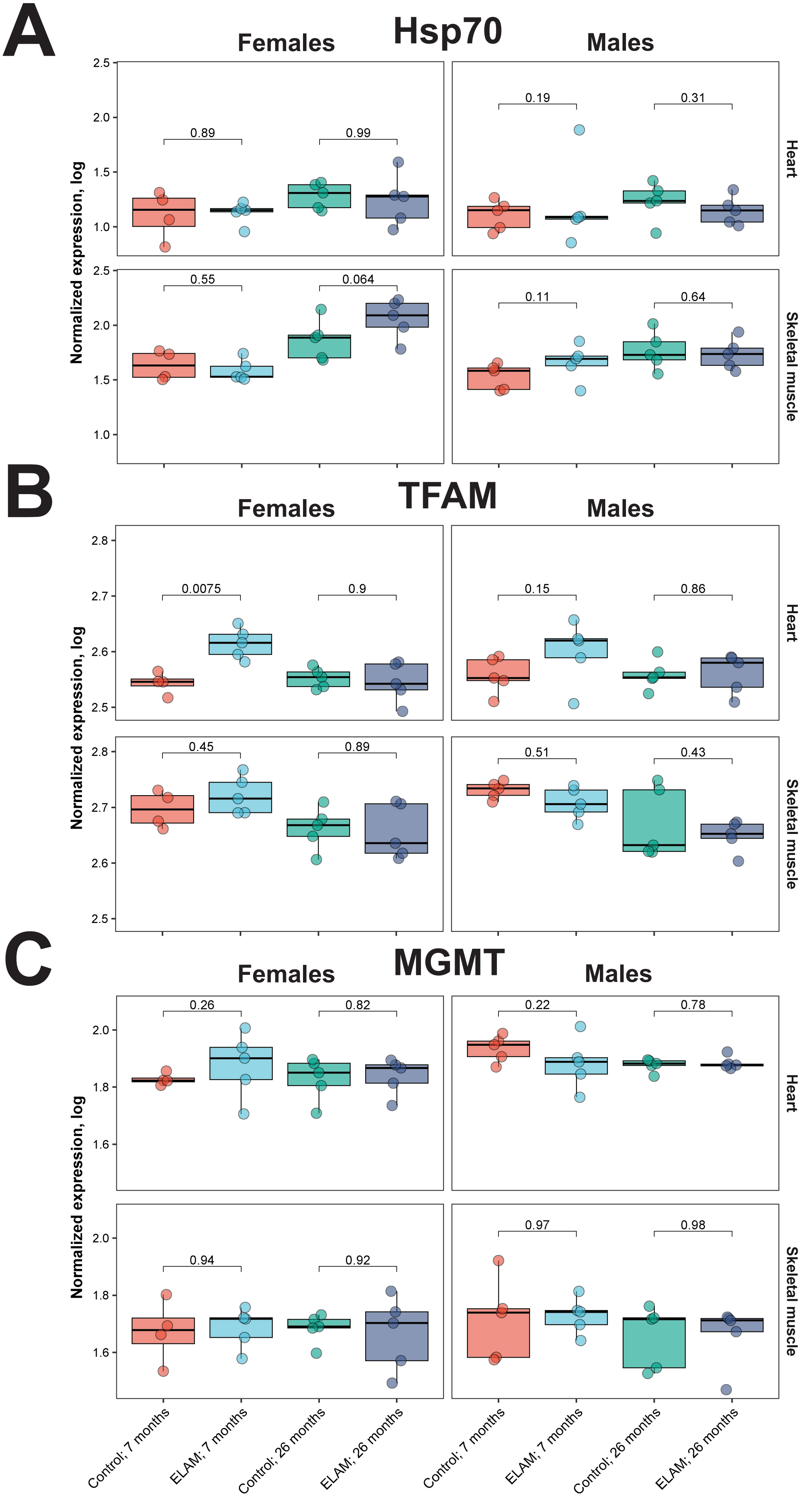


### Extended Data Figure 11
